# Non-junctional Cadherin3 regulates cell migration and contact inhibition of locomotion via domain-dependent opposing regulations of Rac1

**DOI:** 10.1101/750752

**Authors:** Takehiko Ichikawa, Carsten Stuckenholz, Lance A. Davidson

**Affiliations:** Department of Bioengineering, University of Pittsburgh, Pittsburgh, PA, 15260, USA; Nano Life Science Institute (WPI-NanoLSI), Kanazawa University, Kanazawa 920-1192, Japan; Department of Developmental Biology, University of Pittsburgh, Pittsburgh, PA, 15260, USA; Department of Computational and Systems Biology, University of Pittsburgh, Pittsburgh, PA, 15260, USA

**Keywords:** Non-junctional Cadherin, Rac1, Contact inhibition of locomotion, directionality of single cell migration, Xenopus laevis embryo, Gastrulation, Morphogenesis

## Abstract

Classical cadherins are well-known primary adhesion molecules responsible for physically connecting neighboring cells and signaling the cell-cell contact. Recent studies have suggested novel signaling roles for “non-junctional” cadherins (Niessen and Gottardi, 2008; Padmanabhan et al., 2017); however, the function of cadherin signaling independent of cell-cell contacts remains unknown. In this study, we used mesendoderm cells and tissues from gastrula stage *Xenopus laevis* embryos to demonstrate that extracellular and cytoplasmic cadherin domains regulate Rac1 in opposing directions in the absence of cell-cell contacts. Furthermore, we found that non-junctional cadherins regulate contact inhibition of locomotion (CIL) during gastrulation through alterations in the stability of the cytoskeleton. Live FRET imaging of Rac1 activity illustrated how non-junction cadherin3 (formerly C-cadherin) spatio-temporally regulates CIL. Our study provides novel insights into adhesion-independent signaling by cadherin3 and its role in regulating single and collective cell migration *in vivo*.

## Introduction

Cadherins are transmembrane calcium-dependent adhesive molecules that mechanically connect the cytoskeleton of one cell to the cytoskeleton of neighboring cells. Concurrent with this role, cadherins transmit intracellular signals in coordination with other transmembrane proteins, such as epidermal growth factor receptor (EGFR) or P2Y_2_ receptor (P2Y_2_R) that interact through its extracellular domain (Liao et al., 2014; Mateus et al., 2007), or via cytoplasmic proteins such as p120-catenin (p120), β-catenin, and α-catenin through its cytoplasmic domain (Du et al., 2014; Gottardi et al., 2001). These signaling pathways have been studied almost exclusively under conditions where cells maintain cell-cell contacts. Even though recent studies suggest that unbound cadherins, so-called “non-junctional” cadherins (NJCads) can signal through RhoA or β-catenin (Kowalczyk and Reynolds, 2004; Padmanabhan et al., 2017), the functions of NJCads remain elusive.

To investigate the function of NJCads, we adapted assays of single and collective cell migration: (1) contact inhibition of locomotion (CIL) in collectively migrating cells, (2) CIL in single-cell migration, and (3) the directionality of single migrating cells (Becker et al., 2013; Plutoni et al., 2016). CIL was originally reported more than 60 years ago and described the behavior of a cell that stops or changes its direction after colliding with an another, i.e. confronting cell (Abercrombie, 1970; Abercrombie and Heaysman, 1954). Initially, after contact, the leading-edge of the protrusion collapses near the contact site, and a new leading-edge is formed in a different direction (Abercrombie and Heaysman, 1953; Carmona-Fontaine et al., 2008; Mayor and Carmona-Fontaine, 2010). To date, several signal components including RhoA, Rac1, ephrin type-A receptor (EphA), and non-canonical Wnt pathways have been implicated in regulating CIL (Astin et al., 2010; Carmona-Fontaine et al., 2008; Scarpa et al., 2015; Theveneau et al., 2013). Using our quantitative assays, we compared the migration and CIL of single cells and collectively migrating multicellular sheets. To assess collective CIL, we tracked the boundary of migrating mesendoderm sheets and judged whether their forward motion was arrested after the collision with an opposing tissue (Carmona-Fontaine et al., 2008). To assay single-cell CIL, we observed single-cell collisions and quantified the directional change after the collision, as defined in original CIL studies (Abercrombie and Heaysman, 1954).

As junctionally engaged cadherins can confound the role of NJCads, we quantified the directionality of single migratory cells using a standard measure of directionality, expressed as the ratio of the start-to-end distance over the total path-length (Fig. 1H) (George et al., 2013); this ratio is typically less than 1 with lower directionality ratios from less persistent cells. Directionality is known to be regulated by both absolute levels and gradients of Rac1 activation (Pankov et al., 2005; Yamao et al., 2015) where high activity lowers directional migration and low activity reduces directionality (Krause and Gautreau, 2014; Monypenny et al., 2009; Pankov et al., 2005).

**Figure 1.**
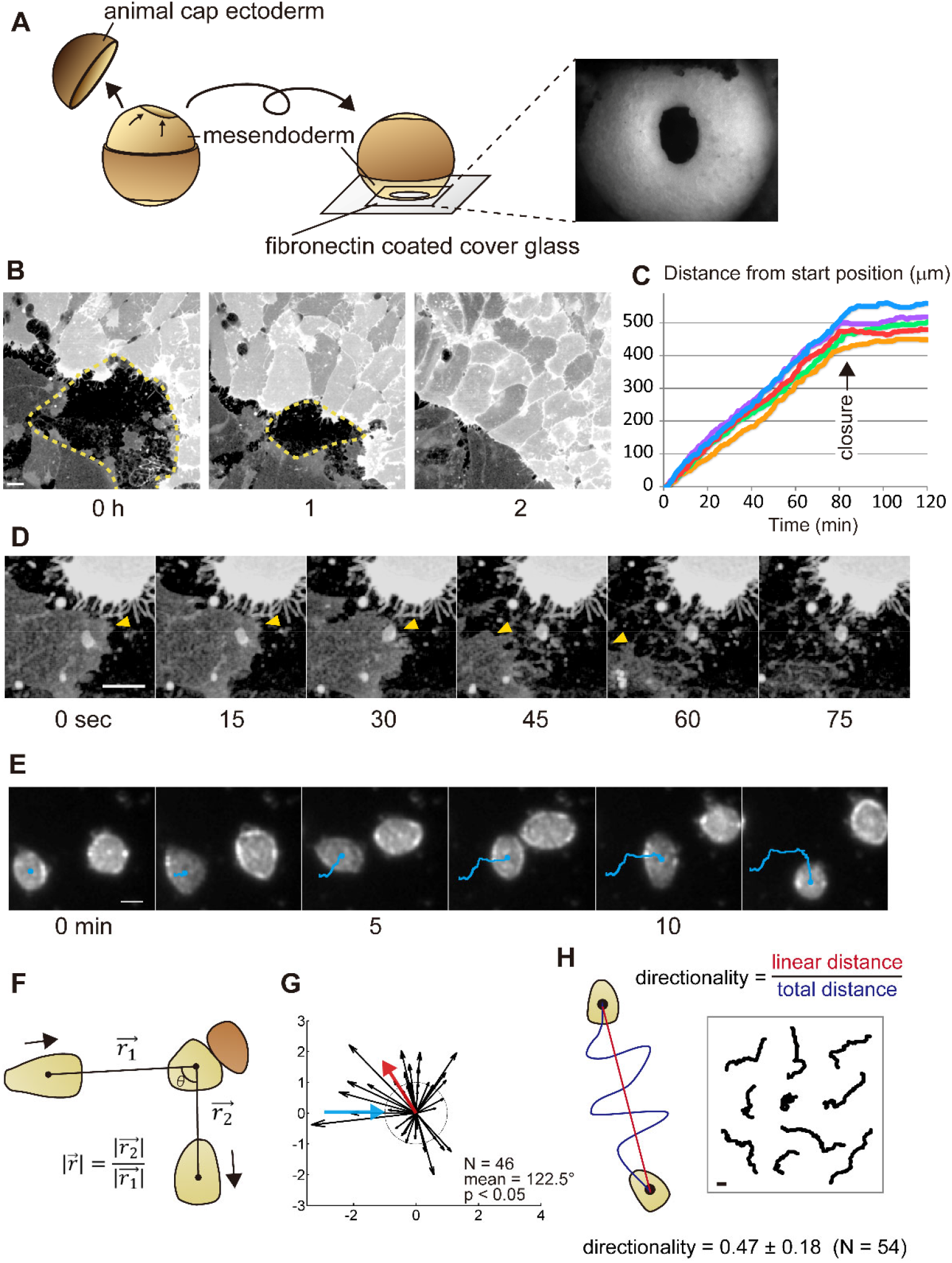
Cell migration assays using *Xenopus* gastrula stage embryonic mesendoderm; contact inhibition of locomotion (CIL) in collective migration, CIL in single migratory cells, and directionality of single motile cells. (A) Schematic of intravital imaging of mesendoderm closure in *Xenopus* embryo from stage 11.5. Animal cap ectoderm was removed, and the lip and outer surface of the mesendodermal mantle was placed in contact with a fibronectin-coated cover glass. The right-side image shows the mesendoderm mantle observed with a stereomicroscope. (B) Frames from a confocal time-lapse showing closing mesendoderm mantle expressing membrane-targeted GFP (dotted lines indicate the boundaries). A difference in the expression level of GFP indicates the different origins of opposing sides. (C) Progressive rates of closure from five embryos. Progress of the leading edge from the start time point to closure at each time points are shown. The arrow indicates the time of the collision. After the collision, cell migration stops. (D) Frames from a representative sequence showing lamellipodia retraction. Leading-edge on the darker cell is indicated by yellow arrowheads; the leading edge lamellipodia retracts after touching the brighter opposing cell. (E) Representative frames from a brightfield time-lapse sequence of colliding single mesendodermal cells. The trajectories are shown with blue lines. WT mesendodermal cells change migration direction after the collision. (F) The geometry of single-cell CIL kinetic analysis with migratory cell (light brown) and opposing cell (dark brown). (G) The vector plot of a set of wild-type (WT) cell collisions (blue arrow - incoming cell; red arrow - mean angle post-collision; dotted circle radius 1 - same velocity before and after collision). For a number of collisions (N), the mean value of the angle taken by the departing cell (incident angle = 0°). Statistical significance is calculated for a circular distribution of post-collision angles (p). Most WT mesendodermal cells change direction after collision indicating CIL. (H) Schematic for measurement of directionality (left) and tracked paths of single migrating cells (right). All scale bars are 20 µm.

Biochemical interactions between cadherins and Rac1 have been previously reported (Goodwin et al., 2003; Kumper and Ridley, 2010; Noren et al., 2001); however, the role of these interactions is not well understood. In *Xenopus*, cadherin3 (cdh3, formerly known as C-cadherin) is the major classical cadherin expressed prior to neurulation and is ubiquitously distributed (Choi and Gumbiner, 1989; Detrick et al., 1990; Winklbauer, 2012). Here, we demonstrate a functional interaction between Rac1 activity and cdh3.

From our motility assays, we found that defects in CIL occur when cells express a mutant form of cdh3 lacking the extracellular domain (ΔE-cdh3). Moreover, expression of ΔE-cdh3 and ΔC-cdh3, a mutant cdh3 that lacks the cytoplasmic domain, produces opposite effects on single-cell directionality and Rac1 activity. For instance, Rac1 activity in cells expressing ΔE-cdh3 is decreased, whereas ΔC-cdh3 expression increases Rac1 activity. The impact of cdh3 mutants on Rac1, CIL, and directionality can be understood through changes in cytoskeletal stability. Furthermore, live-cell imaging of Rac1 activity using a Xenopus-optimized Raichu FRET reporter highlights the spatial and temporal dynamics of Rac1 throughout CIL and how cdh3 mutants disrupt directionality and CIL. Our findings support a model where NJCads, acting through Rac1 and cytoskeletal stability, regulate dynamic changes in cell orientation and persistence, needed for CIL and collective cell movements.

## Results

### Establishment of assay methods for evaluating non-junctional cadherins (NJCads)

To investigate the function of NJCads in mesendoderm during gastrulation, we adapted three assays to quantify motility: CIL in collective migration, CIL in a single cell, and directionality of single-cell migration. To test CIL in collective cell migration, we observed the closure of the mantle-shaped mesendoderm of *Xenopus laevis* embryo at the end of gastrulation stage (Davidson et al., 2002). Mesenchymal mesendodermal cells migrate on the underside of the blastocoel roof toward the animal pole of the embryo (Fig. 1A) to enclose the blastocoel. To confirm that *in vitro* cell movements reproduced *in vivo* behaviors, we used intravital imaging of the mesendoderm mantle (Davidson et al., 2002). Mesendoderm movements can be recorded in minimally manipulated embryos where a portion of the animal cap ectoderm is removed, and the resulting embryo is positioned to place the mesendoderm onto a fibronectin-coated cover glass (Fig. 1A). Mesendoderm cells in these preparations extend large lamellipodia at the leading edge, and the cells elongate as the ring of leading-edge cells converges. CIL can be observed when lamellipodia of leading mesendodermal cells retract after contacting the opposing mesendodermal cells (arrowheads in Fig. 1D, Movie 1). Transient retractions of lamellipodia do not immediately arrest collective migration; however, the collective migration ceased, once mesendodermal cell bodies collided with cell bodies of the opposing margin (Fig. 1B, C). The behavior of migrating sheets of mesendoderm indicates that the mesendoderm of *Xenopus* embryo exhibit CIL similar to CIL observed during collective migration of neural crest *in vivo* (Becker et al., 2013; Carmona-Fontaine et al., 2008; Theveneau et al., 2013).

As junctional cadherins might obscure the role of NJCads, we adapted an assay to quantify CIL in single mesendoderm cells. We dissociated mesendoderm tissue into single cells and tracked single-cell collisions on fibronectin-coated cover glass in time-lapse sequences (Fig. 1E) (Winklbauer and Keller, 1996; Winklbauer and Nagel, 1991). We quantified cell-cell interactions by comparing the change in direction and velocity before and after collision (Fig. 1F) (Carmona-Fontaine et al., 2008; Dunn and Paddock, 1982; Theveneau et al., 2013). We found that wild type (WT) mesendodermal cells significantly change direction after collision (Fig. 1G, Movie 2, and Table S1; N = 46, mean = 122.5°, p < 0.05), indicating that single mesendoderm cells retain the capacity for CIL.

Preliminary tracking of single cells for CIL studies suggested parameters of single-cell motility might also correlate with CIL. To investigate this further, we measured the directionality ratio (d/D, where D is the path length between start and end, and d is the linear distance from start to end) of single migratory cells as they moved 1 hour without cell-cell contact (Fig. 1H). The average directionality of WT mesendodermal cells was 0.47 ± 0.18 (mean ± SE, N = 54).

### Truncation mutant cadherins ΔE-cdh3 and ΔC-cdh3 alter CIL and directional migration in mesendoderm cells

As cdh3 is the major cadherin expressed during gastrulation (Muller et al., 1994), we evaluated CIL after expression of two truncation mutants, ΔE-cdh3 and ΔC-cdh3 (Kurth et al., 1999), lacking extracellular and cytoplasmic domains, respectively. We confirmed the expression of ΔE-cdh3 and ΔC-cdh3 and found that the expression level of these mutants is higher than endogenous cdh3 (Fig. S1). Surprisingly, the mesendoderm sheets expressing ΔE-cdh3 did not stop after collision (Fig. 2A, Movie 3). Instead, leading-edge mesendoderm cells expressing ΔE-cdh3 ran over or invaded opposing mesendoderm in a fashion resembling the behavior of neural crest cells defective in CIL (Carmona-Fontaine et al., 2008; Theveneau et al., 2013) or cell types lacking CIL behaviors (Dunn and Paddock, 1982). However, spreading mesendoderm expressing ΔC-cdh3 alone or ΔE-cdh3 with ΔC-cdh3 exhibited normal CIL and stopped after the collision (Fig. 2A).

**Figure 2.**
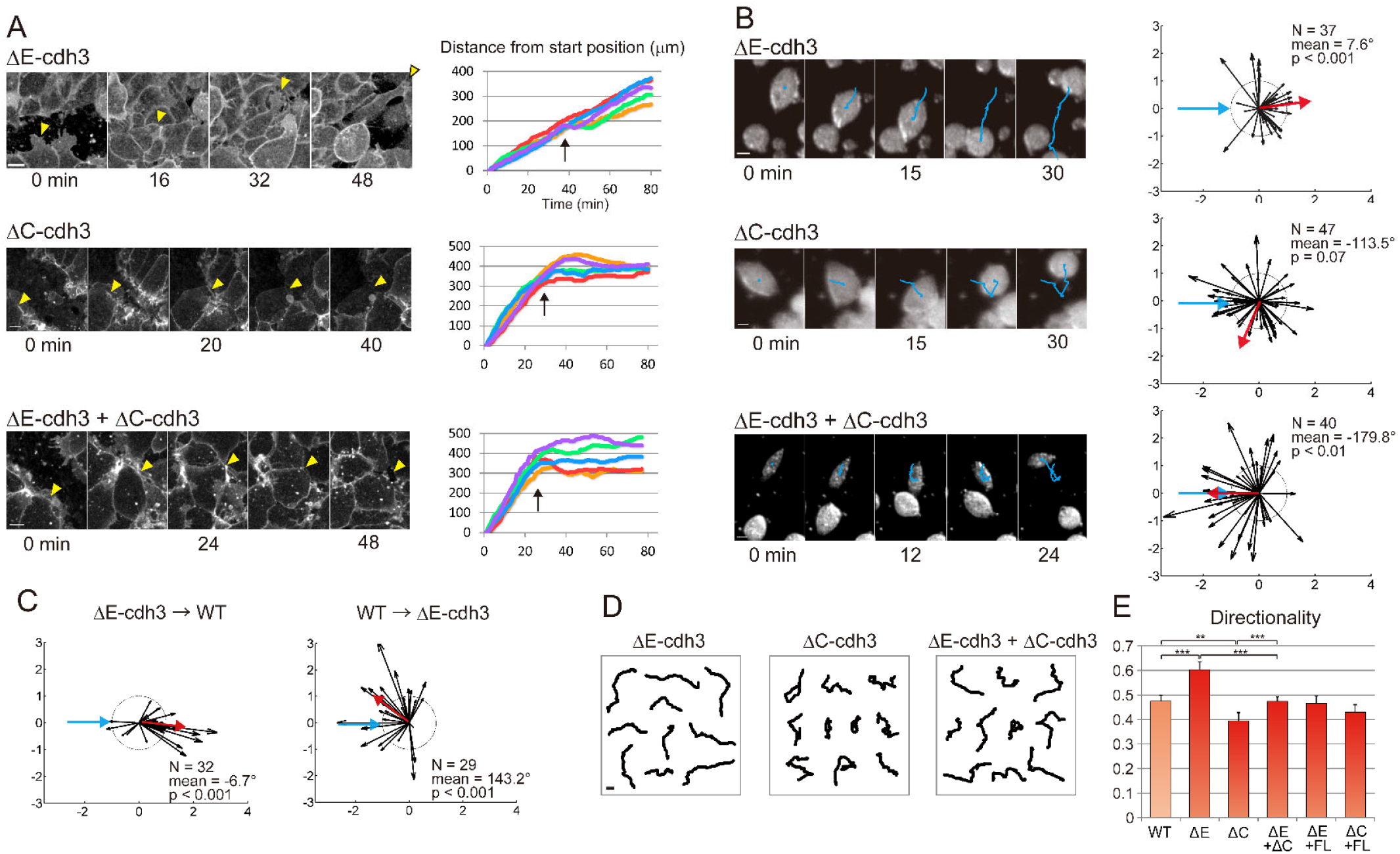
Truncation mutants of cdh3 independently regulate collective CIL and single cell CIL and single cell persistence. (A) Frames from a representative confocal time-lapse sequence (left; leading edge, yellow arrowheads) of membrane-targeted GFP expressing mesendoderm mantle closure in embryos expressing extracellular truncated cdh3 (ΔE-cdh3), cytoplasmic domain truncated cdh3 (ΔC-cdh3), and ΔE-cdh3 + ΔC-cdh3. Positions of leading edge mesendoderm movements during closure (including sequences shown at left, n = 5; arrows indicate time of collision). In ΔE-cdh3 injected embryos, cells continued to migrate after the closure, whereas ΔC-cdh3 and ΔE-cdh3 + ΔC-cdh3 injected embryos stopped after closure. (B) Collisions of ΔE-cdh3, ΔC-cdh3, and ΔE-cdh3 + ΔC-cdh3 expressing single cells (left; trajectories shown in blue). Angle followed by cells after collision (right; mean angle, red; ΔE-cdh3, N = 37, mean = 7.6°, p < 0.001; ΔC-cdh3, N = 40, mean = −179.8°, p < 0.01; ΔE-cdh3 + ΔC-cdh3, N = 47, mean = −113.5°, p = 0.07). ΔE-cdh3 expressing cells did not show CIL, whereas ΔC-cdh3 cells exhibited CIL and ΔE-cdh3 + ΔC-cdh3 injected cells showed partial CIL. (C) Collisions between ΔE-cdh3 expressing cells and WT cells. Left: Summary of collisions of ΔE-cdh3 expressing cells into WT cells (N = 32, mean = −6.7, p < 0.001). The ΔE-cdh3 expressing cells are CIL-defective whereas WT retain CIL. Right: Summary of single cell collisions of WT cells into ΔE-cdh3 expressing cells (N = 29, mean = 143.2, p < 0.001). The WT cells showed CIL. CIL deficiency caused by ΔE-cdh3 is cell autonomous and not the result of loss of cadherin-cadherin binding. (D) Single cell trajectories over 1 hour without collision of ΔE-cdh3, ΔC-cdh3 and ΔE-cdh3 + ΔC-cdh3 expressing cells. (E) Directionality of single cell migration of ΔE-cdh3 (ΔE), ΔC-cdh3 (ΔC), ΔE-cdh3 + ΔC-cdh3 (ΔE + ΔC), ΔE-cdh3 + full length of cdh3 (FL-cdh3) (ΔE + FL), ΔC-cdh3 + FL-cdh3 (ΔC + FL). ΔE-cdh3 expressing cells migrate more persistently than WT or ΔC-cdh3 expressing cells. All scale bars are 20 µm.

Because CIL could be junction-dependent, we carried out single-cell collisions between mesendodermal cells expressing ΔE-cdh3 and found that they lacked CIL (Fig. 2B, Movie 4; N = 37, mean = 7.6°, p < 0.001); whereas cells expressing ΔC-cdh3 showed CIL. Co-expression of ΔE-cdh3 and ΔC-cdh3 partially rescued the deficiency of CIL caused by expression of ΔE-cdh3 alone (Fig. 2B, Movie 5, 6; N = 47, mean = −113.5°, p = 0.07). Deficiency of CIL by expressing ΔE-cdh3 was also partially rescued by co-expressing full-length cdh3 (FL-cdh3, Fig. S3A). Importantly, we confirmed that CIL was cell-autonomous, as CIL was retained when WT cells encountered ΔE-cdh3 cells but were CIL-deficient when ΔE-cdh3 expressing cells encountered WT cells (Fig. 2C). Furthermore, we found that junctions between extracellular domains of cdh3 were not responsible for CIL because cells did not exhibit CIL in collisions with cdh3 conjugated beads (Fig. S2). Thus, our data suggest that CIL can be regulated via NJCads signaling.

To eliminate the possibility of junction effect of cadherins, we further assessed the effects of cdh3 mutants on single-cell directionality without cell-cell contact, a measure of migratory persistence (Pankov et al., 2005; Petrie et al., 2009; Winklbauer et al., 1992). ΔE-cdh3 expressing cells were 27% more directional than WT cells (Fig. 2D, E, and Table S2; in contrast to WT trajectories shown in Fig. 1H). Conversely, ΔC-cdh3 expressing cells exhibited 17% lower directionality than WT. Directionality was also restored to wild-type levels when full-length cdh3 was co-expressed with either ΔE-cdh3 and ΔC-cdh3. These results demonstrate that the extracellular and cytoplasmic domains of cdh3 have the opposite functions on persistence, a key factor in cell motility; and the intracellular and extracellular cdh3 domains might activate and inhibit, respectively, the same target pathway.

### Expression of ΔE-cdh3 or ΔC-cdh3 decreases or increases the Rac1 activity

The effect of mutant cadherins on single-cell persistence is reminiscent of the effects of Rac1 (Matthews et al., 2008; Pankov et al., 2005) and suggests that Rac1 is inhibited in ΔE-cdh3 expressing cells and activated in ΔC-cdh3 expressing cells. To confirm, we measured the activity of Rac1 in mesendoderm tissues isolated from ΔE-cdh3, ΔC-cdh3, and ΔE-cdh3 and ΔC-cdh3 co-expressing embryos (Hara et al., 2013; Zhang et al., 1998). Rac1 activity in ΔE-cdh3 expressing tissues was approximately 0.4-fold of WT levels, and Rac1 activity of ΔC-cdh3 expressing tissues was approximately 1.8-fold higher than WT (Fig. 3A, B, and Fig. S6). In tissues co-expressing ΔE-cdh3 and ΔC-cdh3, the activity was almost equal as in WT tissues. Thus, the extracellular or the cytoplasmic domains of cdh3, respectively, activate or inhibit Rac1 activity in migratory mesendoderm.

**Figure 3.**
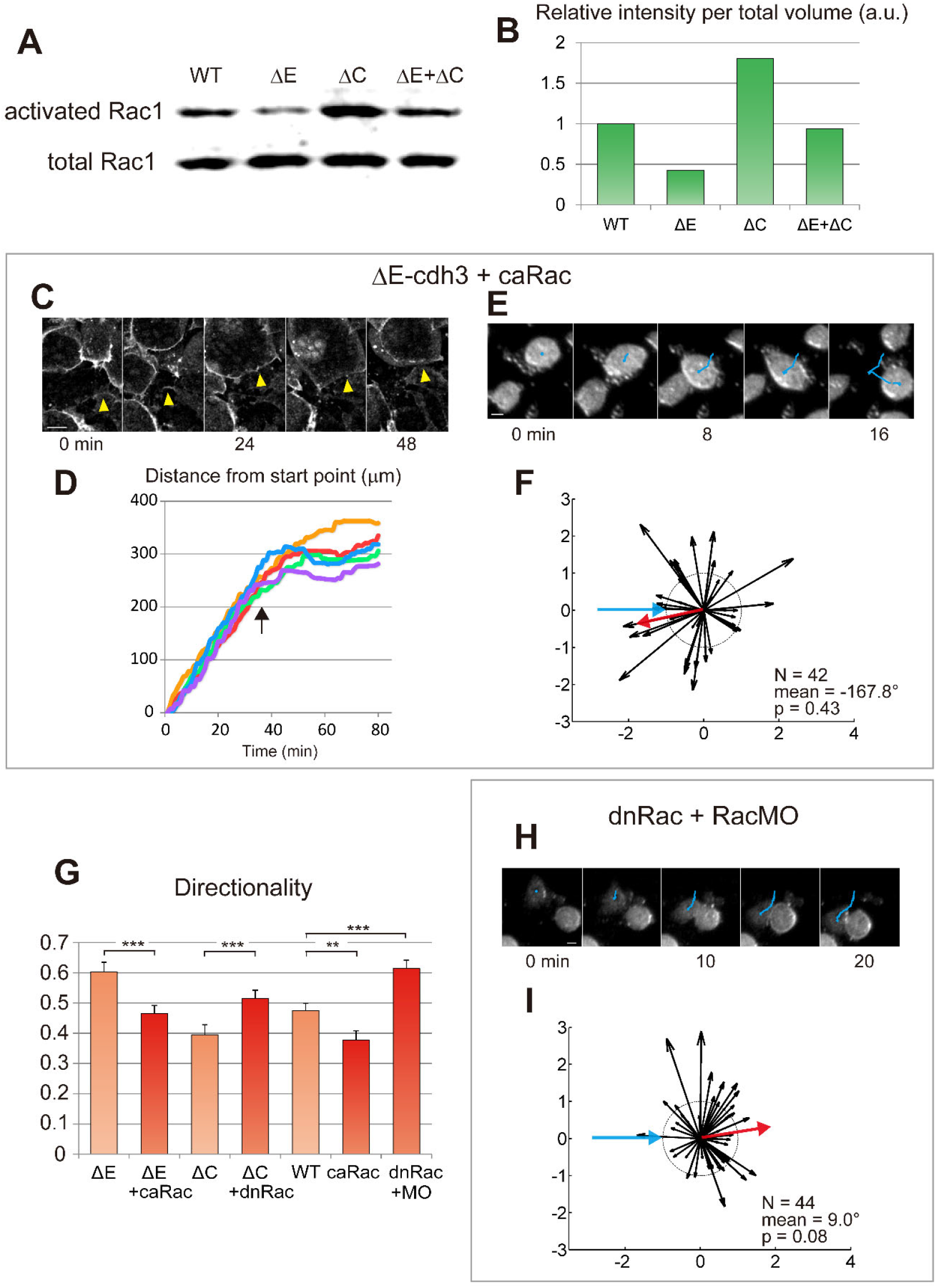
Truncated cadherins control Rac1 activity responsible for single cell CIL, collective CIL, and single cell persistence. (A) Rac1 activity is modulated from WT by ΔE-cdh3 (ΔE), ΔC-cdh3 (ΔC), and ΔE-cdh3 + ΔC-cdh3 (ΔE + ΔC). (B) Rac1 activity of (A) normalized to WT levels. Rac1 activity in ΔE-cdh3 and ΔC-cdh3 expressing cells was 0.4- and 1.8-fold of WT levels. ΔE + ΔC activity was not significantly different from WT levels. (C) Mesendoderm closure in embryos co-injected with ΔE-cdh3 + constitutively active form of Rac1 (caRac). (D) Time-courses of mesendoderm closure from five embryos. CIL-defects caused by ΔE-cdh3 were rescued by co-injection of caRac. (E) Single cell collisions of ΔE-cdh3 + caRac expressing cells. (F) Summary of collisions of ΔE-cdh3 + caRac expressing cells (N = 42, mean = −167.8°, p = 0.43). CIL-defects caused by ΔE-cdh3 were partially rescued. (G) Directionality of single cell migrations of ΔE-cdh3 + caRac (ΔE + caRac), ΔC-cdh3 + dominant negative form of Rac1 (dnRac) (ΔC + dnRac), caRac and dnRac + Rac1 morpholino oligomers (RacMO) (dnRac + RacMO) compared with ΔE-cdh3 (ΔE), ΔC-cdh3 (ΔC) and WT, respectively (light red). (H) Single cell collisions of dnRac + RacMO co-injected cells. (I) Summary of collisions of dnRac + RacMO co-injected cells (N = 44, mean = 9.0°, p = 0.08). dnRac + RacMO cells showed the partial defects in CIL. All scale bars are 20 µm.

To directly measure the interaction of cdh3 and rac1, we assessed whether Rac1 activation or inhibition could rescue the CIL and cell motility changes induced by ΔE-cdh3 or ΔC-cdh3. We co-injected ΔE-cdh3 with a constitutively active form of Rac1 (caRac) (Wittmann et al., 2003) and found that Rac1 restored collective CIL, and furthermore, restored CIL in single-cell assays. The mesendoderm movement after closure and the defects in single-cell CIL caused by ΔE-cdh3 were partially rescued by co-expression with caRac1 (Fig. 3C-F; N = 42, mean = −167.8°, p = 0.43). Co-injection of ΔE-cdh3 with caRac and co-injection of ΔC-cdh3 with a dominant-negative form of Rac1 (dnRac) (Habas et al., 2003) rescued both the high and low directionality of single migratory cells caused by ΔE-cdh3 and ΔC-cdh3, respectively (Fig. 3G and Table S2). Similarly, mild inhibition of Rac1 using a small molecule inhibitor (20 nM NSC23766; (Gao et al., 2004)) also rescued the low directionality of ΔC-cdh3 expressing cells (Fig. S3B and C). These results indicate that changes in CIL and motility induced by ΔE-cdh3 and ΔC-cdh3 can be rescued by compensatory modulation of Rac1 activity.

We also tested whether modulated Rac1 activity alone could alter CIL and directionality. In order to reduce Rac1 activity as low as possible, we co-injected dnRac with the antisense morpholino against endogenous Rac1 (RacMO, Fig. S4). The co-injection of dnRac and RacMO reduced Rac1 activity to 0.07 fold of WT (Fig. S4A, B). The inhibition of Rac1 activity by the co-injection of dnRac with RacMO showed partial inhibition of CIL and increased directionality (Fig. 3G-I; N = 44, mean = 9, p = 0.08). High concentrations of Rac1 inhibitor (0.3 mM NSC23766) showed similar results when applied to WT cells (Fig. S3C; N = 49, mean = −11.5°, p = 0.17). caRac injection also resulted in low directionality (Fig. 3G). These results support a mechanism where cadherin controls CIL and cell migration by regulating Rac1 activity.

### Rac1 activated cell boundary domains are reduced in ΔE-cdh3 expressing cells and increased in ΔC-cdh3 expressing cells

To investigate the spatial and temporal dynamics of Rac1 activity during cell-cell collision in ΔE-cdh3 and ΔC-cdh3 expressing cells, we adapted the Raichu-Rac FRET biosensor (Itoh et al., 2002; Kardash et al., 2010) for use in *Xenopus*, and confirmed that the FRET signal reflects dynamic Rac1 activity in *Xenopus* mesendodermal cells (Fig. S5). To analyze the polarity of Rac1 activity, we quantified FRET signal at cell membranes using custom image analysis code (Fig. 4A-C) in WT, ΔE-cdh3, and ΔC-cdh3 expressing cells during cell-cell collisions. In WT cells, Rac1 activity was high at the front of the migrating cell as reported previously (Fig. 4D-F) (Kraynov et al., 2000; Machacek et al., 2009). During CIL, Rac1 activity at the contact site disappeared within two minutes and increased at a different location, and following contact, the cell migrated in the direction of the new site of Rac1 activity (Fig. 4D, E, and Movie 7). Furthermore, contacts initiated at sites of high Rac1 induced stronger CIL than contacts first occurring at sites of low Rac1 (Fig. 4M). In ΔE-cdh3 expressing cells, Rac1 activity was lower throughout the cell and did not change during collisions (Fig. 4G-I, and Movie 8), and the cells continued to migrate in the same direction as prior to the collision. In contrast, in ΔC-cdh3 expressing cells, Rac1 was activated at multiple locations along the cell periphery, and migration ceased in the initial direction after collision (Fig. 4J-L, and Movie 9). To quantify the spatial pattern of high Rac1 activity along the cell periphery, we calculated the percentage of the cell’s boundary where FRET signal ratio was higher than 1.2 for the cell. We found high Rac1 activity over 12.6 ± 0.18% of the periphery of WT cells (N = 57), 7.6 ± 0.22% of ΔE-cdh3 (N = 50, p < 0.01), and 27.3 ± 0.26% of ΔC-cdh3 (N = 59, p < 0.001). Live analysis of Rac1 activity revealed that CIL coincided with the transfer of Rac1 activity away from sites of contact and that spatial domains of high Rac1 activity were further restricted in ΔE-cdh3 and expanded in ΔC-cdh3 expressing cells.

**Figure 4.**
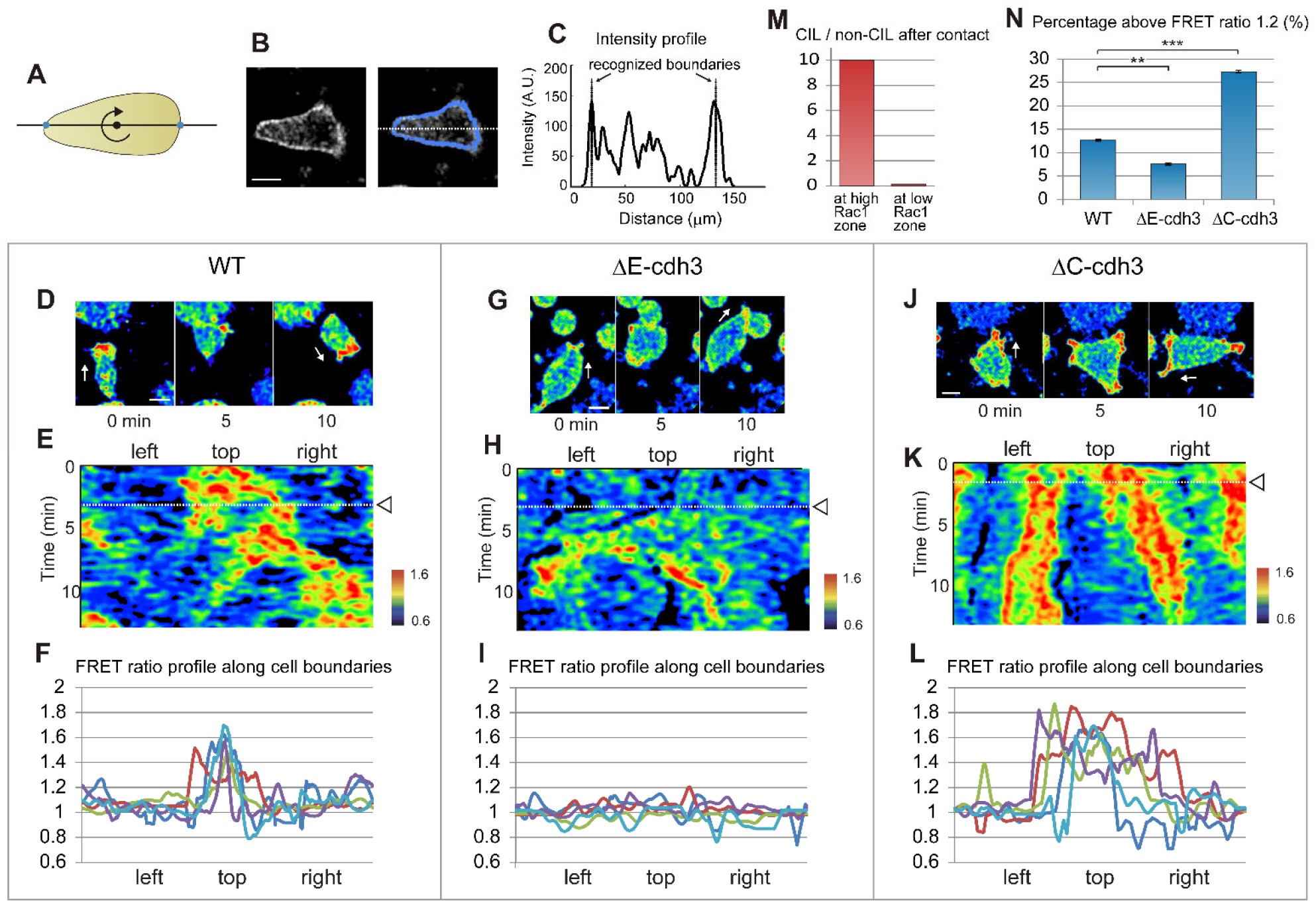
Truncated cadherins modulate the spatial patterns of Rac1 activity during cell-cell collisions. (A) Schematic method segmenting cell membrane. The line passing through the center of cell detects peaks on both sides in the YFP channel and rotates 360° by 1°. (B) Representative Rac1 biosensor FRET image (CFP channel, left) and segmented points for activity quantification (blue line, right). (C) The intensity profile of Rac1 activity along the white dotted line (B; cell boundary indicated). (D) Representative FRET images before, during, and after a typical cell-cell collision of single WT cell. Rac1 activity is shown in pseudocolor; white arrows indicate the directions of cell migrations. (E) Kymograph of Rac1 activity along the cell perimeter during a cell-cell collision with cell initially moving to the top of the frame in (D). The center of x-axis indicates the front of the migrating cell prior to the collision. The arrowhead and dotted line indicate the time of the collision. After the collision, Rac1 activity at the contact position is quickly reduced and reappears at a different location as Rac1 activity increases. (F) FRET profile along cell perimeters of five typical cells. The center of x-axis indicates the front of the migrating cell. (G) Time-course images before, during, and after a typical cell-cell collision of single ΔE-cdh3 expressing cell. (H) Kymograph during the cell-cell collision of ΔE-cdh3 expressing cell. Rac1 activity is low in the whole-cell, and there is little evidence of active Rac1 or polarized activity. (I) FRET ratio profile along cell perimeters of five typical cells. (J) Time-course images before, during, and after a typical cell-cell collision of single ΔC-cdh3 expressing cell. (K) Kymograph during the cell-cell collision of ΔC-cdh3 expressing cell. Rac1 activity was high at multiple positions around the cell periphery. (L) FRET ratio profile along cell perimeters of five typical cells. (M) Frequency of WT CIL is high when the first contact occurs in a high Rac1 activity zone and low when contact occurs in a low Rac1 activity zone. (N = 77 contacting at high Rac1 zone, 9 contacting at low Rac1 zone). (N) Percentage of cell perimeter above ratio 1.2 along cell boundary in WT, ΔE-cdh3, and ΔC-cdh3 cells. WT: 12.6 ± 0.18 (mean ± SE)% (N = 57), ΔE-cdh3: 7.6 ± 0.22% (N = 50, p < 0.01), ΔC-cdh3: 27.3 ± 0.26% (N = 59, p < 0.001).

### NJCads regulates cell migration through regulating cytoskeleton stability

Domain dependent NJCad signaling through Rac1 activity might regulate CIL and cell directionality by modulating microtubule or F-actin dynamics (Kaverina and Straube, 2011; Wittmann et al., 2003). Such a role for Rac1 implies that defects in CIL in ΔE-cdh3 expressing cells are due to a destabilized cytoskeleton. To validate this hypothesis, we modulated cytoskeletal stability in ΔE-cdh3 expressing cells, increasing the stability of microtubules with paclitaxel (microtubule stabilizer) and F-actin with jasplakinolide (F-actin stabilizer). Low doses of either cytoskeletal stabilizer partially rescued the defect in CIL and reduced high directionality caused by ΔE-cdh3 (Fig. 5A, B, and E; N = 49, mean = −25°, p = 0.71, b; N = 37, mean = −22.9°, p = 0.09 and for directionality E, and Table S2). Conversely, we observed that low doses of both nocodazole, to destabilize microtubules, and cytochalasin D, to destabilize F-actin, both lowered CIL and increased directionality of WT cells (Fig. 5C, D and E; N = 38, mean = 30.7°, p = 0.19, d; N = 42, mean = 7.1°, p = 0.87 and for directionality E and Table S2). Even though further studies are needed to identify the precise molecular mechanisms coupling Rac1 to cdh3, our results reveal an NJCad role for ΔE-cdh3 as it perturbs CIL through Rac1, either destabilizing microtubules or F-actin.

**Figure 5.**
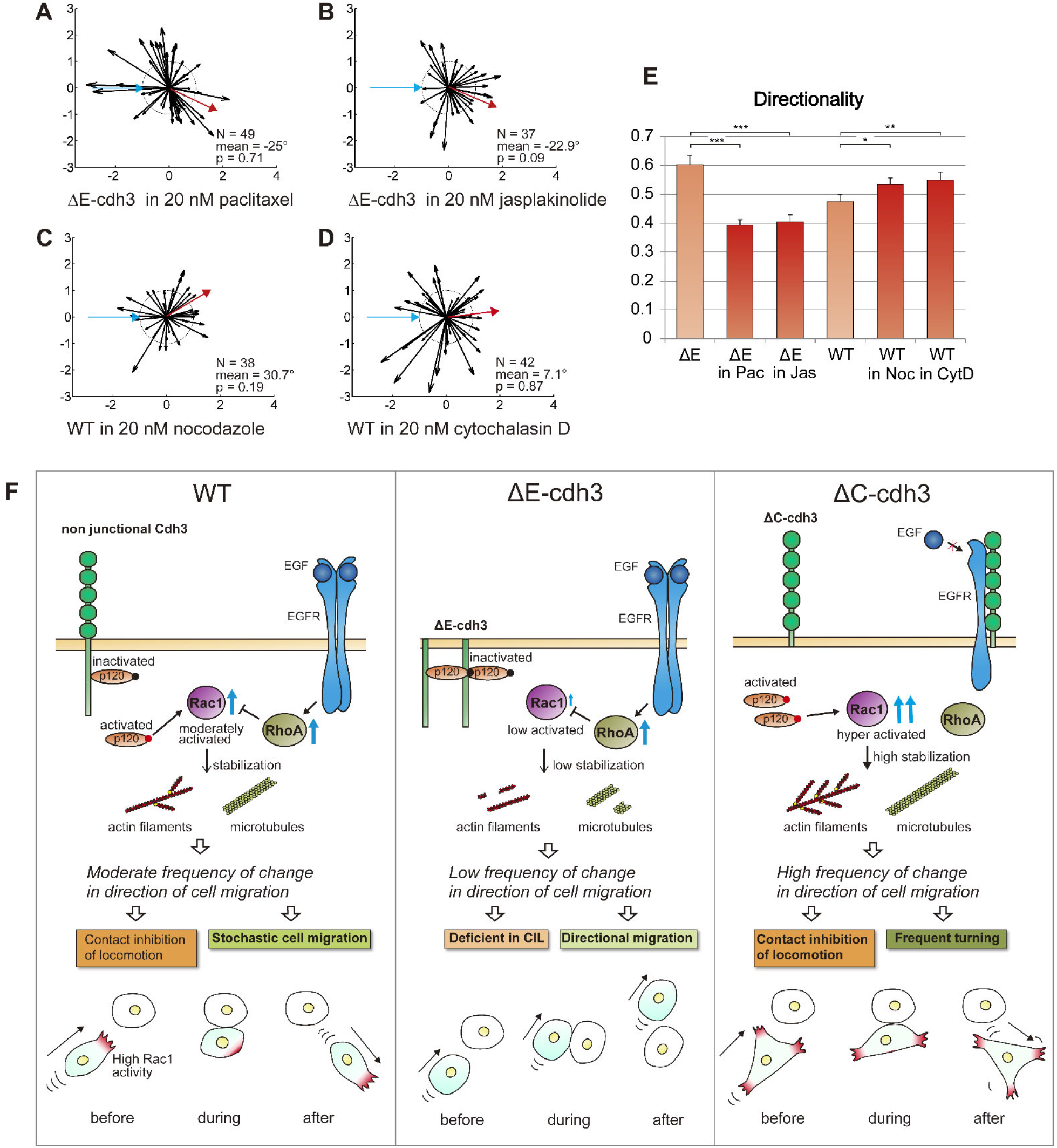
Effects of non-junctional cadherin3 on single-cell CIL and persistence of single-cell migration mitigated by changes in cytoskeletal stability. (A) Summary of collisions of ΔE-cdh3 expressing cells in 20 nM paclitaxel (N = 49, mean = −25, p = 0.71). This mild stabilization of microtubule polymerization partially rescued CIL defects caused by ΔE-cdh3. (B) Summary of collisions of ΔE-cdh3 expressing cells in 20 nM jasplakinolide (N = 37, mean = −22.9°, p = 0.09). The mild stabilization of f-actin polymerization also partially rescued the CIL defects caused by ΔE-cdh3. (C) Summary of collisions of WT cells in 20 nM nocodazole (N = 38, mean = 30.7°, p = 0.19). The mild inhibition of microtubule polymerization caused partial defects in CIL. (D) Summary of collisions of WT cells in 20 nM cytochalasin D (N = 42, mean = 7.1°, p = 0.87). The mild inhibition of f-actin polymerization also caused partial defects in CIL. (E) The quantified directionality of single-cell migration of ΔE-cdh3 cell in 20 nM Paclitaxel (ΔE in Pac), ΔE-cdh3 cell in 20 nM Jasplakinolide (ΔE in Jas), WT cell in 20 nM Nocodazole (WT in Noc), and WT cell in 20 nM Cytochalasin D (WT in CytD) compared with ΔE-cdh3, and WT, respectively (light red). (F) Model of the hypothetical mechanism regulating CIL and directionality of single-cell migration in cells expressing non-junctional cdh3 or truncated cdh3s through localized Rac1 activity and cytoskeleton stability. Rac1 is regulated independently by extracellular and cytoplasmic domains of non-junctional cadherin oppositely through p120 and/or EGFR. In WT cells, Rac1 activity is held at moderate levels by the extracellular and cytoplasmic domains of non-junctional cdh3. Moderate Rac1 activity leads to moderate levels and gradient of cytoskeletal stability. As a stable cytoskeleton supports persistent migration, the moderate stability of cytoskeleton in WT cells results in the moderate frequency of change in the direction of cell migration and enables CIL and moderate directionality of single-cell migration. In ΔE-cdh3 expressing cell, expression of the cytoplasmic domain of cdh3 increases inactivated p120, suppressing Rac1 activity and destabilizing the cytoskeleton; resulting in a low frequency of change in the direction of cell migration, defective-CIL, and lowered persistence. Meanwhile, ΔC-cdh3 suppresses RhoA, which inhibits Rac1 through EGFR and simultaneously fails to deactivate p120, and highly activates Rac1 and leads to the stable cytoskeleton and high frequency of change in the direction of cell migration, resulting in frequent turning and cells capable of CIL.

## Discussion

Non-junctional cdh3 signaling differentially regulates the Rac1 pathway through extracellular and cytoplasmic domains and can modulate CIL and the directionality of single-cell migration through cytoskeletal stability. Based on our manipulations, we propose a model in which the non-junctional extracellular and the cytoplasmic domain of cdh3 controls Rac1 activity in opposite ways, altering cytoskeletal stability to regulate CIL and the persistence of cell migration (Fig. 5F). Stability of the cytoskeleton determines the frequency of change in the direction of cell migration; this results in the difference of the response for CIL and cell migration directionality among WT and truncated cadherins. Rac1 is well known for its role in driving persistent cell migration and collective migration as well as its ability to modulate the stability of the cytoskeleton (Petrie et al., 2009); however, its regulation by non-junctional cdh3 is surprising.

How might non-junctional cadherin3 regulate Rac1? Studies on cadherin1 (formerly known as E-cadherin) implicate the cytoplasmic factor p120 in activating both Rac and Rho (Fang et al., 2004; Franz and Ridley, 2004; Valls et al., 2012). p120 is thought to be active when it is in the cytoplasm and inhibited when it binds to the membrane-proximal cytoplasmic domain of cadherin (Anastasiadis and Reynolds, 2001; Scarpa et al., 2015). Expressed ΔE-cdh3 might inactivate p120 by binding to the cytoplasmic domain of ΔE-cdh3, whereas expressed ΔC-cdh3 might repress the expression level of endogenous cdh3 (Troxell et al., 1999), and the freely available p120 might activate Rac1 (Fig. 5F). Cadherin1 (cdh1; formerly E-cadherin) can also modulate signaling through the EGFR when the extracellular cdh1 blocks binding of EGF to EGFR, preventing activation of RhoA (Mateus et al., 2007). Under these conditions, Rac1 may be indirectly activated by the release of Rac1 inhibition by RhoA (Rottner et al., 1999). Alternatively, the intact extracellular domain of ΔC-cdh3 might inhibit ligand binding to EGFR, leading to increase Rac1 activity (Fig. 5F).

Extracellular- or cytoplasmic-domain deleted mutants of cadherin have been a principal tool in understanding the role of cell-cell adhesion during development. At high levels, ΔE-cdh2 (formerly known as N-cadherin) in *Xenopus* embryo induces lesions in the ectoderm at the end of gastrulation (Kintner, 1992). ΔE-cdh2 in zebrafish embryo led to non-directional migration of facial branchiomotor neurons and defect in caudal tangential migration (Rebman et al., 2016). Similar results have been reported after overexpression of cdh1 mutants (Levine et al., 1994). Overexpression of ΔC-cdh3 in *Xenopus* embryo induces a delay in blastopore closure and defect of posterior patterning, leaving intact anterior region (Lee and Gumbiner, 1995). In our experiments, ΔE-cdh3 overexpression also produces lesions in the ectoderm and delay of the blastopore closure (not shown). We also confirmed the earlier observation that ΔC-cdh3 overexpression delays blastopore closure. In both cases, the mesendoderm mantle retains the ability to close by at the end of gastrulation. It is uncertain whether non-junctional cadherin signaling contributes to these phenotypes; however, our findings suggest reinvestigation of the role of Rac1 in these lesions. Kashef’s group demonstrated that expression of cytoplasmic-domain truncated cdh1, ΔC-cdh1, rescued defects in neural crest migration after cdh1 knockdown, and that embryos expressing the extracellular domain deleted mutant, ΔE-cdh1, showed severe migration defects (Huang et al., 2016). These defects have been commonly attributed to defects in cell-cell adhesion, but our findings suggest another possibility, for instance, that ΔC-cdh1 might rescue cell motility defects by restoring Rac1 activity and that ΔE-cdh1 may inhibit neural crest motility by inhibiting Rac1.

Our study demonstrates that cdh3 has non-junctional roles in collective migration and raises the question of how non-junctional signaling integrates with junctional signaling. Cadherin engagement is known to regulate RhoA and Rac1 activity (Liu et al., 2006; Noren et al., 2003; Noren et al., 2001; Perez et al., 2008). We presume some cadherins remain non-junctional even after cell-cell contact is established. We propose that during collective migration, both junctional and non-junctional cadherin signals work simultaneously; however, further studies are expected to clarify this mechanism.

Cohesotaxis, a process wherein pulling force using adhesive beads touched to cell surface, transmits a signal to the cell to move to the opposite direction of applied force, maybe one of the cases in which junctional and non-junctional cadherin signals work simultaneously (Roca-Cusachs et al., 2013; Weber et al., 2012). In this case, cadherin apparently works as an adhesive molecule; however, we would propose a broader role for cadherin3 that includes Rac1 signaling in mechanosensing by NJCads, as they regulate Rac1 activity and cytoskeleton stability.

## Materials and methods

### *Xenopus laevis* embryos and constructs

*Xenopus laevis* embryos were obtained by *in vitro* fertilization using standard protocols (Sive et al., 2000). Fertilized eggs were de-jellied in 2% cysteine solution (pH 8) and were microinjected at the one/two-cell stage at multiple sites with a total of 0.5 ng (for FRET biosensors) or 1 ng (for other mRNAs) of mRNAs in 1× modified Barths Solution (MBS) containing 4% Ficoll-400 (17-0300-05, GE Healthcare) using a microinjector (PLI-90, Harvard Apparatus) (Kim and Davidson, 2011). Embryos were transferred to 1/3× MBS and cultured until mid-gastrula (st. 11.5).

Deletion mutants of the extracellular or cytoplasmic domain of cadherin3, ΔE-cdh3, and ΔC-cdh3 were constructed from the full length of C-cadherin (Lee and Gumbiner, 1995) and were kind gifts from T. Kurth and P. Hausen (Kurth et al., 1999). cdh3 (cdh3-a), formerly known as C-cadherin, is the only classical cadherin expressed during gastrulation, and E/N-cadherin is not expressed during this stage. We adopted the name cadherin3 instead of C-cadherin, to specify cdh3-a in our study, and not cdh3-b, another variant, also known by the name C-cadherin. Our Rac1 morpholino antisense oligonucleotide (RacMO; Gene Tools) was designed against the translational initiation site of *Xenopus laevis* Rac1 (5’-CCACACATTTAATGGCCTGCATGGC-3’). The effectiveness of the RacMO mediated knock-down was confirmed by western blot and showed that 12 ng total injections reduced Rac1 protein levels (Fig. S4A, C) (Habas et al., 2003).

### Imaging conditions and quantitative methods of collective and single-cell migration

To assess single and collective cell movements including CIL, we recorded live-cell movements using confocal microscopy methods, previously established for tracking *Xenopus* gastrula stage mesendodermal cell, and explants expressing membrane-targeted GFP (Davidson et al., 2002). For intravital, explant, and single-cell preparations, we first microsurgically removed a large patch of ectoderm from the animal pole region of a stage 11.5 embryo, exposing cells at the margin of the mesendoderm mantle. For intravital imaging, we placed the mesendoderm mantle and remainder of the embryo onto a fibronectin-coated glass coverslip (Fig. 1B). This configuration allowed the mesendoderm mantle, including leading-edge cells, contact with fibronectin along with the same interfaces as they would encounter *in vivo* (Davidson et al., 2002; Davidson et al., 2004). In cases where the mesendoderm mantle closure is limited (e.g., ΔE-cdh3), we removed two pieces of mesendoderm and placed them on the fibronectin-coated glass, so the leading edges of the two pieces would spread and make contact (Davidson et al., 2002). Confocal time-lapse sequences were acquired at 15-second intervals with a high N.A. 63× oil immersion objective using an inverted confocal microscope (TCS SP5, Leica). Migrating cells were tracked, and their motion quantified using the Manual Tracker plug-in of ImageJ (http://rsb.info.nih.gov/ij/plugins/track/track.html) and analyzed with Excel (2010, Microsoft). To quantify mesendoderm closure, we calculated the distance between the rim at the start point and at each time point. Time-to-close graphs were drawn to align individual trajectories from different experiments to the same closure time.

To assay single-cell migration and contact dynamics, we recorded the movement of mesendoderm cells dissociated from mesendoderm mantles described above. Dorsal mesendodermal cells were isolated at stage 11.5 in Ca^2+^-Mg^2+^-free DFA (53 mM NaCl, 5 mM Na_2_CO_3_, 4.5 mM K Gluconate, 32 mM Na Gluconate) and transferred into low Ca^2+^-Mg^2+^-DFA (0.2 mM Ca^2+^ and 0.2 mM Mg^2+^) in a small glass-bottomed chamber coated with fibronectin. Low Ca^2+^ and Mg^2+^ media allowed mesendoderm cells to migrate and collide but not aggregate as they would in standard media containing 1 mM Ca^2+^ and 1 mM Mg^2+^. Varying the concentration of Ca^2+^ and Mg^2+^ in single and collective cell experiments did not affect the directionality of single-cell migration (data not shown). Cell movements were recorded at 15-second intervals with a bright-field inverted microscope (Axiovert S100, Carl Zeiss) equipped with a digital camera (CFW-1308M, Scion Corporation). The following inhibitors were used: NSC23766 (733767-34-5, Santa Cruz Biotechnology), paclitaxel (T7191, Sigma-Aldrich), jasplakinolide (42017, Calbiochem), nocodazole (487928, Calbiochem), and cytochalasin D (C8273, Sigma-Aldrich).

CIL in single cells was quantified by established methods (Carmona-Fontaine et al., 2008; Dunn and Paddock, 1982; Theveneau et al., 2013). Each cell collision was tracked with Manual Tracker of ImageJ plug-in. The location of cell-cell contacts and cell positions, 5 minutes prior to-and after-contact were analyzed. CIL parameters of angle and the ratio of the velocity before and after the collision were calculated and plotted using MATLAB (2012a, Mathworks). Statistical analysis performed with Excel calculated the mean angle and circular p-value (Rayleigh test (Zar, 1999); Table S1). The mean angle was measured with respect to the incident angle of the approaching cell. Normal CIL was defined in collisions, where the mean angle was significantly greater than 120° (p < 0.05), and defective-CIL was defined by collisions, where the mean angle was significantly less than 60° (p < 0.05). To quantify directionality, we tracked single cells that migrated without contacting other cells for one hour, using the Manual Tracking plug-in of ImageJ. Cell directionalities (start-to-end distance divided by path length) were calculated with Excel.

CIL in single cells encountering cadherin coated beads used cdh3-coated beads prepared by binding purified cdh3-fc (gifted from Dr. Douglas W. DeSimone and Dr. Barry. M. Gumbiner) to Protein A/G coated beads (53132, Thermo Fisher Scientific) (Chappuis-Flament et al., 2001). Briefly, 2-fold excesses C-cadherin-fc needed to saturate the beads were incubated in 10 μM HEPES, 50 mM NaCl, pH 7.2, 1 mM CaCl_2_ for 90 minutes at 4°C on an Eppendorf shaker (1,400 rpm). Cdh3-coated beads were pelleted, washed twice, and resuspended in 400 μl of 10 mM HEPES, and 50 mM NaCl, pH 7.2. Cadherin conjugation was confirmed by western blot using anti-cdh3 antibody (6B6, Developmental Studies Hybridoma Bank) and Odyssey CLx Imaging System (Fig. 1K, 9140, LI-COR). Polyacrylamide beads (1504150, BIO-RAD) were used as controls.

### Rac1 activity and western blotting

Bulk Rac1 activity was measured using an assay kit (BK035, Cytoskeleton), as previously described (Hara et al., 2013). Embryos were dissected and broken up in lysis buffer (50 mM Tris pH 7.5, 10 mM MgCl_2_, 0.5 M NaCl and 2% Igepal) with 1/100 (v/v) protease inhibitor cocktail (contained in the kit), 1/100 (v/v) phosphatase inhibitor cocktail 2 (P6726, Sigma Aldrich), 1/100 (v/v) phosphatase inhibitor cocktail 3 (P0044, Sigma Aldrich), and 25 mM sodium fluoride (919, Sigma Aldrich) to inhibit GTP hydrolysis (Vincent et al., 1998). The lysates were centrifuged at 14,000 g for 10 minutes x 5 times to remove lipid. The supernatant was used in a pull-down assay with PAK-PBD beads that bind active Rac1, and the products were run on a conventional protein blot. In brief, products were denatured by the addition of the same volume of 2× SDS sample buffer (0.5 M Tris–HCl; pH 6.8, 10% (w/v) SDS, 50% (w/v) glycerin, 5% (v/v) 2-mercaptoethanol). Samples were then boiled for 5 minutes and processed by SDS-PAGE, blotted onto a PVDF membrane (162-0177, BIO-RAD), reacted with the 1:1000 anti-Rac1 monoclonal antibody (610651, BD Transduction Laboratories), and detected using 1:5000 IRDye 800CW anti-mouse antibody (926-32351, LI-COR) and Odyssey CLx Imaging System (9140, LI-COR). The expression level of the cdh3 and cdh3 mutant proteins was measured in the same protocol of western blot used for measuring Rac1 activity. Protein extracts were as described previously (Baronsky et al., 2016). After transfer, blots were probed with mouse anti-cdh3 (C-cadherin; Clone 6B6, Developmental Studies Hybridoma Bank, dilution 1:300), mouse anti-cdh1 (E-cadherin; Clone 5D3, Developmental Studies Hybridoma Bank, dilution 1:300), anti-Myc (Clone 9E10, Sigma-Aldrich, dilution 1:1,000), and rabbit anti-ɣTubulin (T3320, Sigma-Aldrich, dilution 1:3,000). Secondary antibodies were IRDye 680LT Goat anti-Mouse IgG (Licor 925-68020) and IRDye 800CW Goat anti-Rabbit IgG (Licor 925-32211). Anti-myc was used to detect FL-cdh3, ΔC-cdh3, and ΔE-cdh3. Anti-cdh3 was used for FL-cdh3 and ΔC-cdh3. Anti-tubulin was used for loading controls.

### Imaging subcellular dynamics of Rac1 using FRET probe and the analysis

To visualize Rac1 activity in living cells, we adapted a Raichu-Rac FRET probe (gift from Dr. Erez Raz (Kardash et al., 2010)). However, the original FRET sensor did not localize to the plasma membrane in our cells. To improve membrane association, we replaced the C-terminal CAAX domain with one from the H-Ras C-terminal domain, which facilitates membrane localization in *Xenopus* (Fig. S5). The modified Raichu-Rac probe showed clear localization and significantly enhanced FRET signal at the membrane and reduced non-specific activity in the cytoplasm (Fig. S5). Time-lapse sequences of the FRET signal at 15-second intervals were acquired from an inverted confocal microscope (TCS SP5, Leica) using spectral detection of the YFP channel (535 ± 15 nm) and CFP channel (480 ± 20 nm) with excitation from the 458 nm argon laser line.

The FRET signal was calculated using previously published protocols (Kardash et al., 2010). In brief, YFP and CFP channels of the Raichu-Rac1 FRET signal were individually processed by smoothing and background corrected. The FRET ratio image ([YFP]/[CFP]) was calculated using ImageJ. The signal intensities at the cell boundaries were calculated using custom MATLAB scripts at 360 points with 10-point moving average (Fig. 4A-C, codes are available upon request).

The number of CIL events (in Fig. 4M) was counted when the direction of cell migration significantly changed within 5 minutes after the contact. Rac1 activity along a cell perimeter was categorized as “high” when the FRET ratio was greater than 1.2 and “low” when the ratio was less than 1.2.

## Acknowledgments

We thank T. Kurth and P. Hausen for providing the constructs of extracellular (ΔE-cdh3) and cytoplasmic (ΔC-cdh3) domain deficient of cdh3-a (formerly called Cadherin-C), R. Habas for the constructs of constitutively active and negative form of Rac1, E. Raz for Raichu-Rac FRET biosensor and N. Hukriede for the use of Odyssey imaging systems. C-cadherin-fc was gifted by D. W. DeSimone and B. M. Gumbiner. We thank Y. Hara, A. Shindo and T. S. Yamamoto for advice on quantifying Rac1 activity and Davidson lab members for providing support with the experiments and revising the manuscript. This study was supported by The Uehara Memorial Foundation (to T.I.) and National Institutes of Health grants (NIH) (R01 HD044750 and R21 ES019259) and the National Science Foundation (NSF) (CAREER IOS-0845775 and CMMI-1100515). Any opinions, findings, and conclusions or recommendations expressed in this material are those of the authors and do not necessarily reflect the views of the NSF or the NIH.

## Competing interest statement

The authors declare no competing financial interests.

## Supplementary material

Supplementary material for this article is available at http://dev.biologists.org/lookup/suppl/doi:

## Supplementary materials

**Figure S1.**
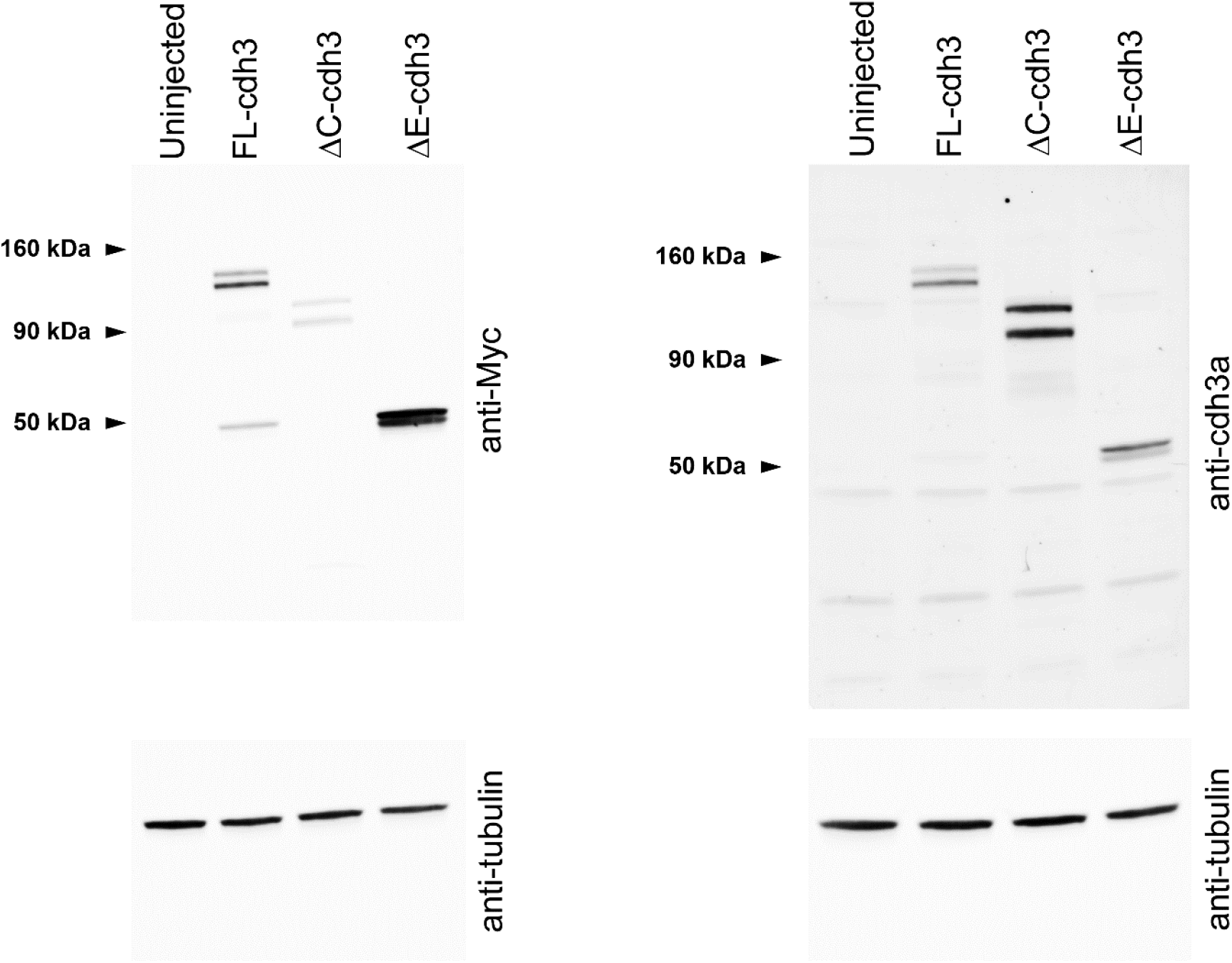
The expression level of FL-cdh3, ΔE-cdh3, and ΔC-cdh3. Western blot analysis of uninjected, FL-cdh3, ΔE-cdh3, and ΔC-cdh3 injected embryos using anti-Myc or anti-cdh3a compared with tubulin. Several isoforms (precursor and processed) are shown (Baronsky et al., 2016; Choi and Gumbiner, 1989).

**Figure S2.**
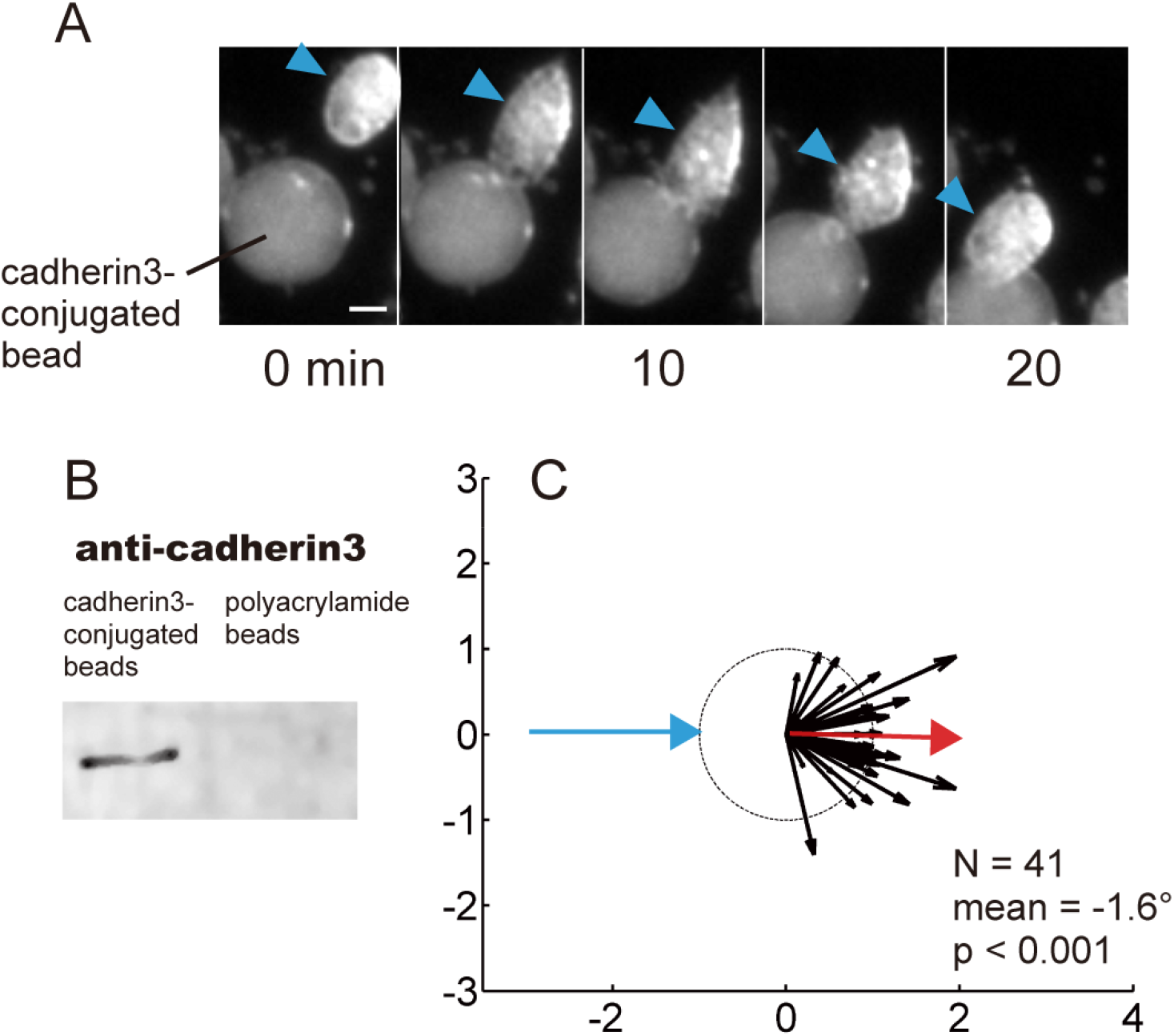
Collisions to cadherin3 conjugated beads. (A) Time sequence of the collision between a WT mesendodermal cell and a cadherin3-conjugated bead. (B) Western blot analysis of cadherin3-conjugated beads (Protein A/G beads) and control beads (polyacrylamide beads). (C) Summary of collisions between WT cells and cadherin3-conjugated beads (N = 37, mean = 7.6°, p < 0.001). Collisions between cells and cdh3-beads did not result in CIL. Scale bar: 20 μm.

**Figure S3.**
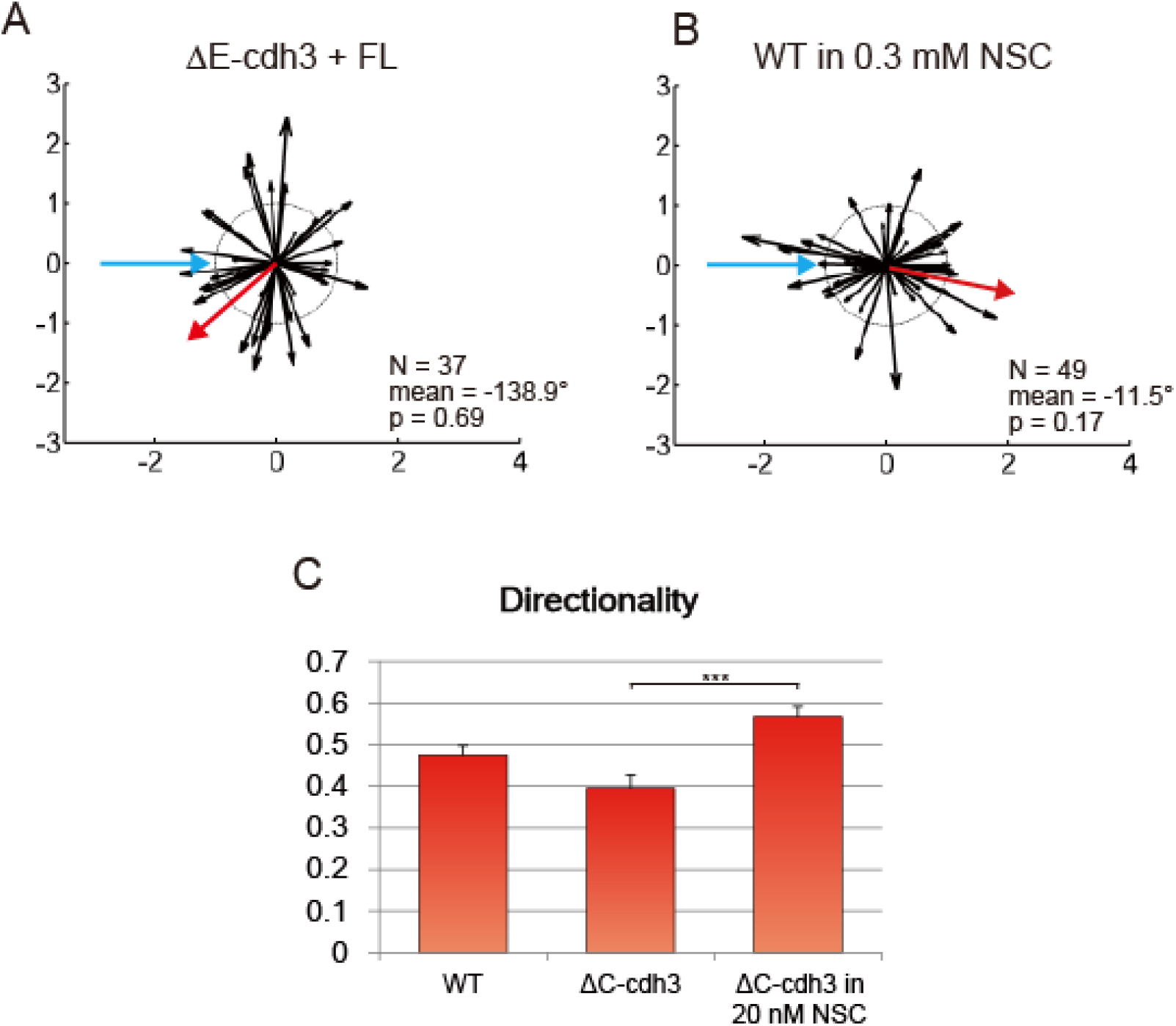
Single-cell CIL and single-cell persistence under various conditions. (A) Summary of collisions of ΔE-cdh3 + FL-cdh3 (N = 37, mean = −138.9, p = 0.69). CIL defects caused by ΔE-cdh3 are partially rescued by co-expression of FL-cdh3. (B) Summary of collisions of WT in 0.3 mM NSC23766 (N = 49, mean = −11.5, p = 0.17). The inhibition of Rac1 activity induces partial defects in CIL. (C) Low persistence of ΔC-cdh3 expressing cells is enhanced by Rac1 inhibition by 20 nM NSC23766.

**Figure S4.**
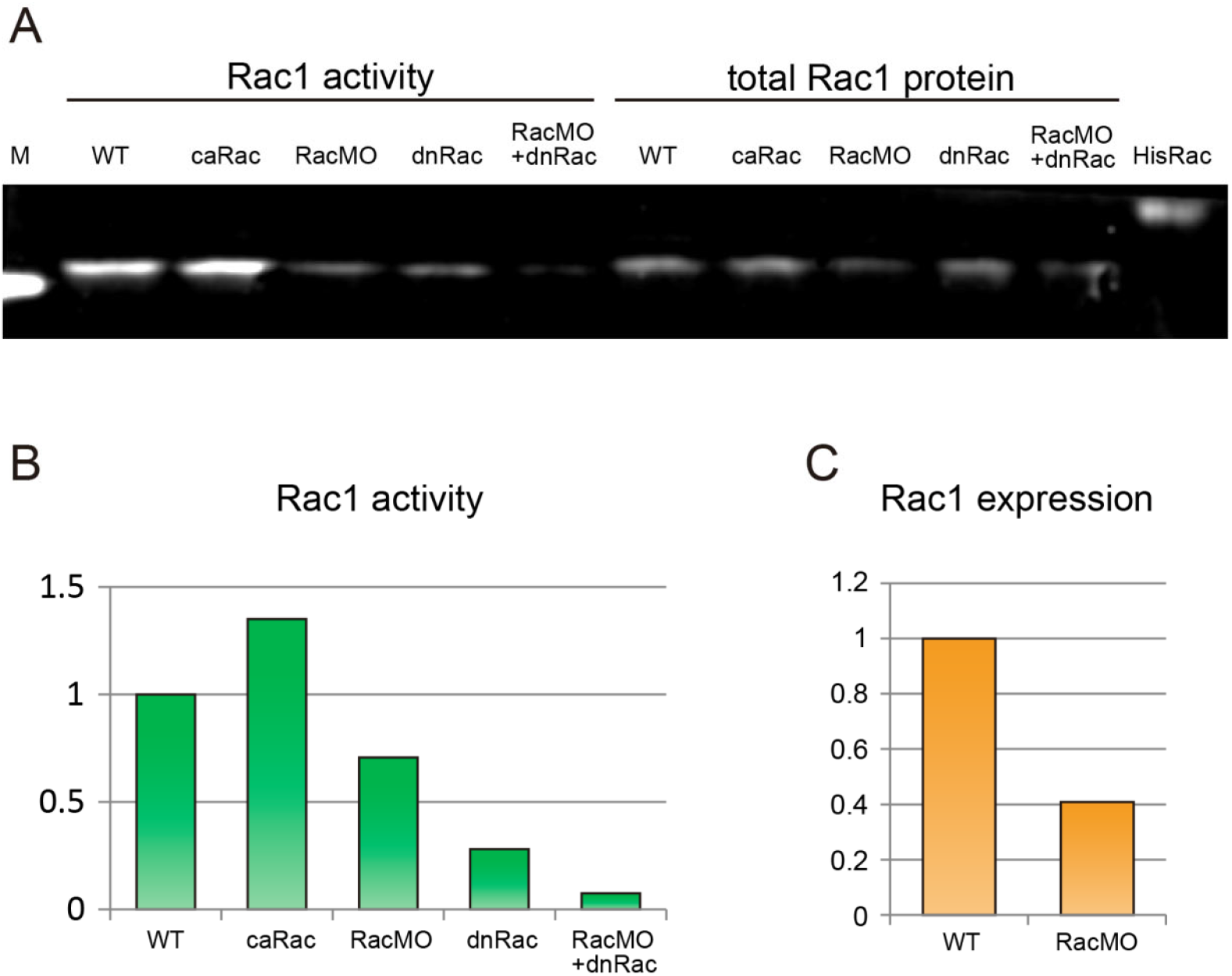
CaRac, RacMO, and dnRac alter Rac1 activity; RacMO reduces Rac1 abundance and activity. (A) Blot showing Rac1 activity and total Rac1 expression. M: marker band of 20 kDa, WT: wild type control, HisRac: His-tag conjugated purified Rac1 protein (24 kDa). (B) Quantification of Rac1 activation from (A) normalized to WT activity. The relative Rac1 activity after expression of caRac, RacMO, dnRac and RacMO + dnRac was 1.34, 0.7, 0.28, and 0.07 respectively. (C) The comparison of Rac1 protein expression between WT and RacMO from (A) normalized to WT expression levels. 12 ng injection of RacMO reduced Rac1 protein expression to 40% of WT levels.

**Figure S5.**
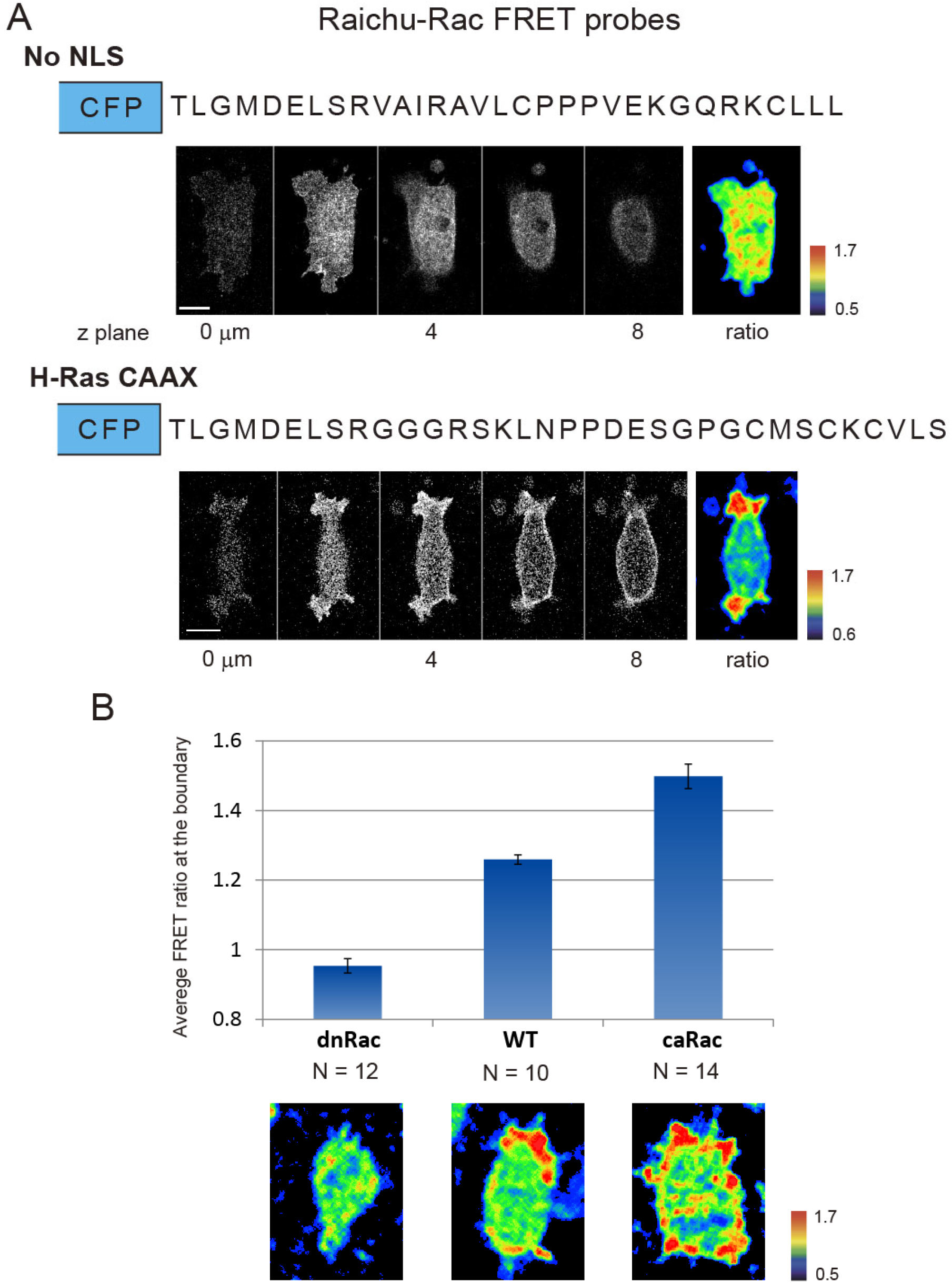
Modified Raichu-Rac FRET biosensor with improved membrane localization. (A) Comparison between Raichu-Rac without the nuclear localization signal (no NLS)(Kardash et al., 2010) and new Raichu-Rac with C-terminal domain of H-Ras (H-Ras CAAX). The top panel shows the amino acid sequences of C-terminal domain, slices from a z-series confocal stack of the YFP channel and the ratio of no NLS. The bottom panel shows FRET signal from H-Ras CAAX modified biosensor and clear polarity of Rac1 activity. (B) The response of Raichu-Rac with H-Ras CAAX in cells expressing dnRac and caRac. The FRET ratio value was calculated by averaging the FRET signal along the cell periphery over 10 minutes.

**Figure S6.**
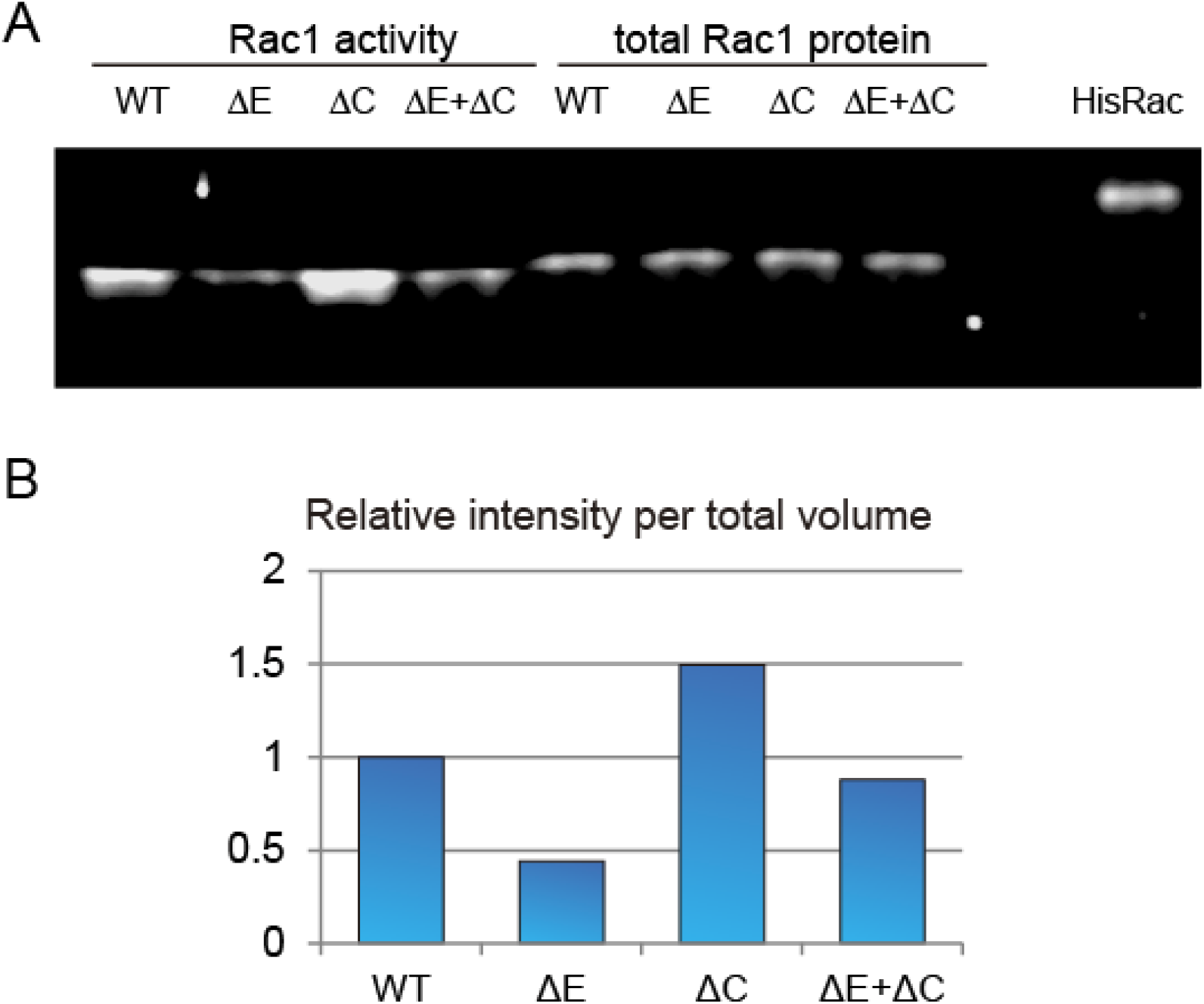
An additional blot of Rac1 activation of wild type and mutant cadherin injected embryo shown in. Fig. 3A **and** B. (A) Rac1 activation analysis in WT, ΔE-cdh3 (ΔE), ΔC-cdh3 (ΔC), and ΔE-cdh3 + ΔC-cdh3 (ΔE + ΔC) injected embryos. (B) Rac1 activity of (A) normalized to WT levels.

**Movie 1.**
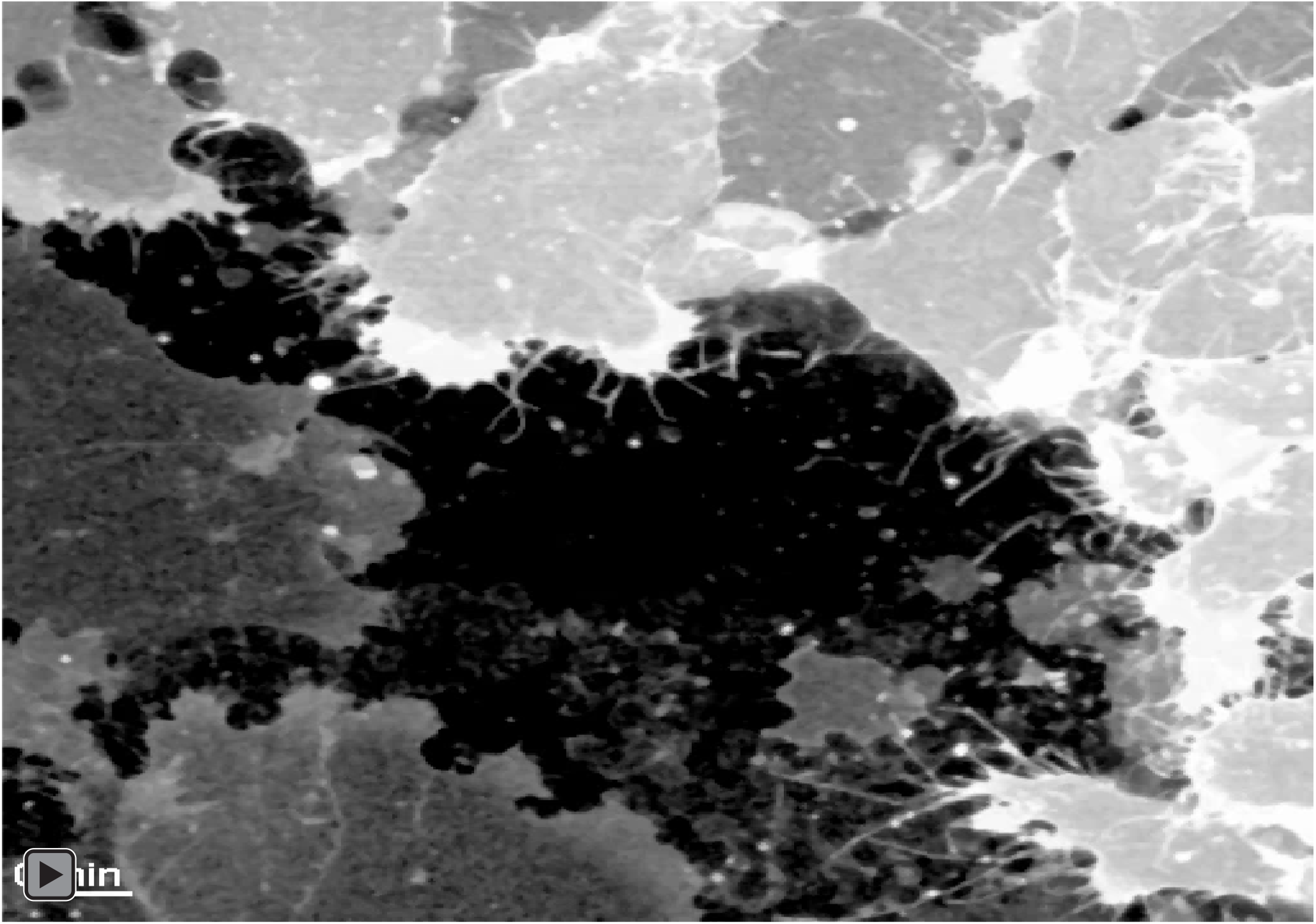
Mesendoderm closure within wild type embryo. Intravital confocal time-lapse sequence of mesendoderm closure expressing membrane-targeted GFP. The different expression level of GFP indicates the cells originating from different regions around the marginal zone. Images were acquired every 15 seconds with a high N.A. oil-immersion 63× lens and an inverted confocal microscope. Scale bar is 20 µm.

**Movie 2.**
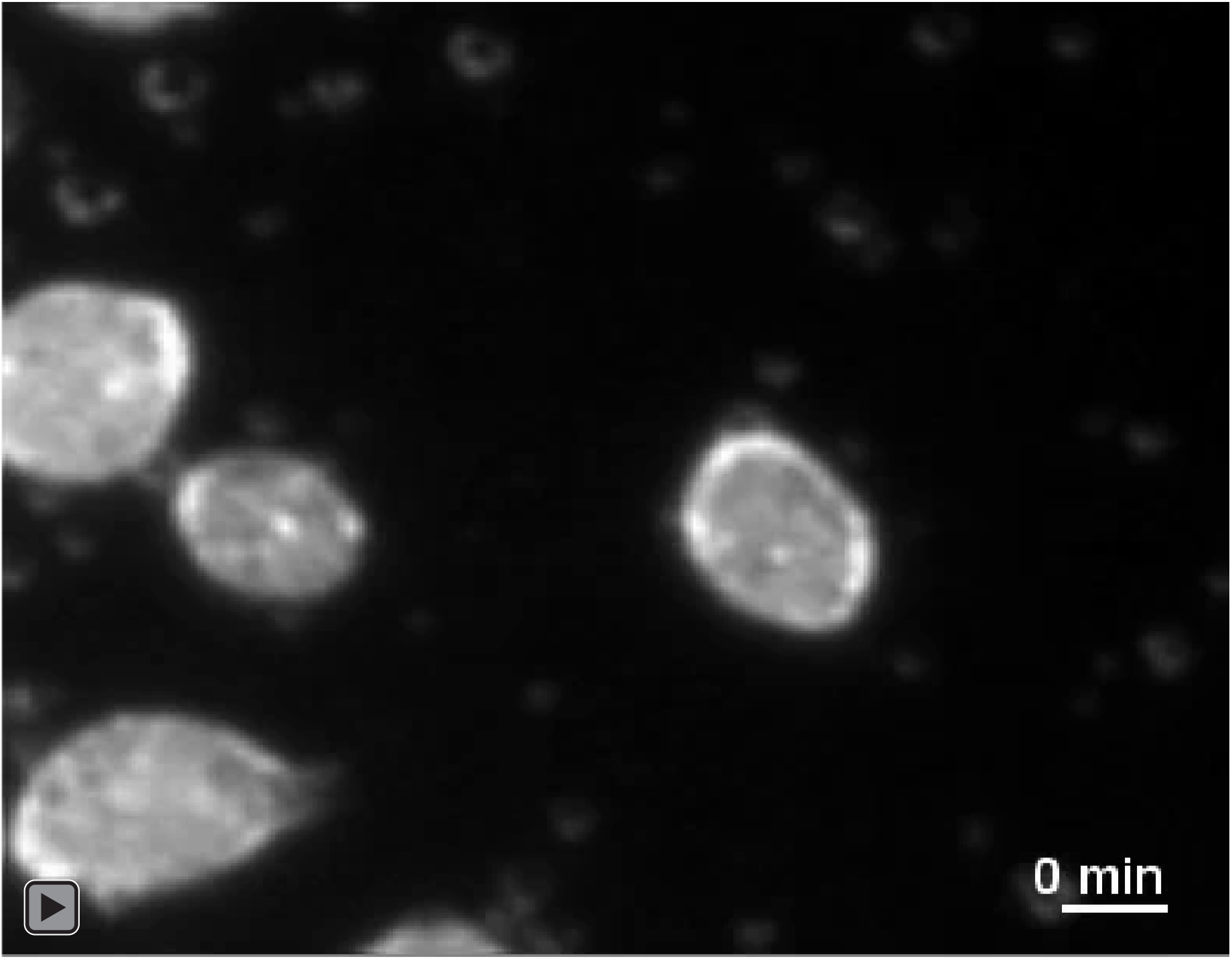
Single-cell CIL between wild type mesendoderm cells. The time-lapse sequence shows CIL between single wild type mesendodermal cells observed using brightfield illumination on an inverted compound microscope. The time interval is 30 seconds. Scale bar is 20 µm.

**Movie 3.**
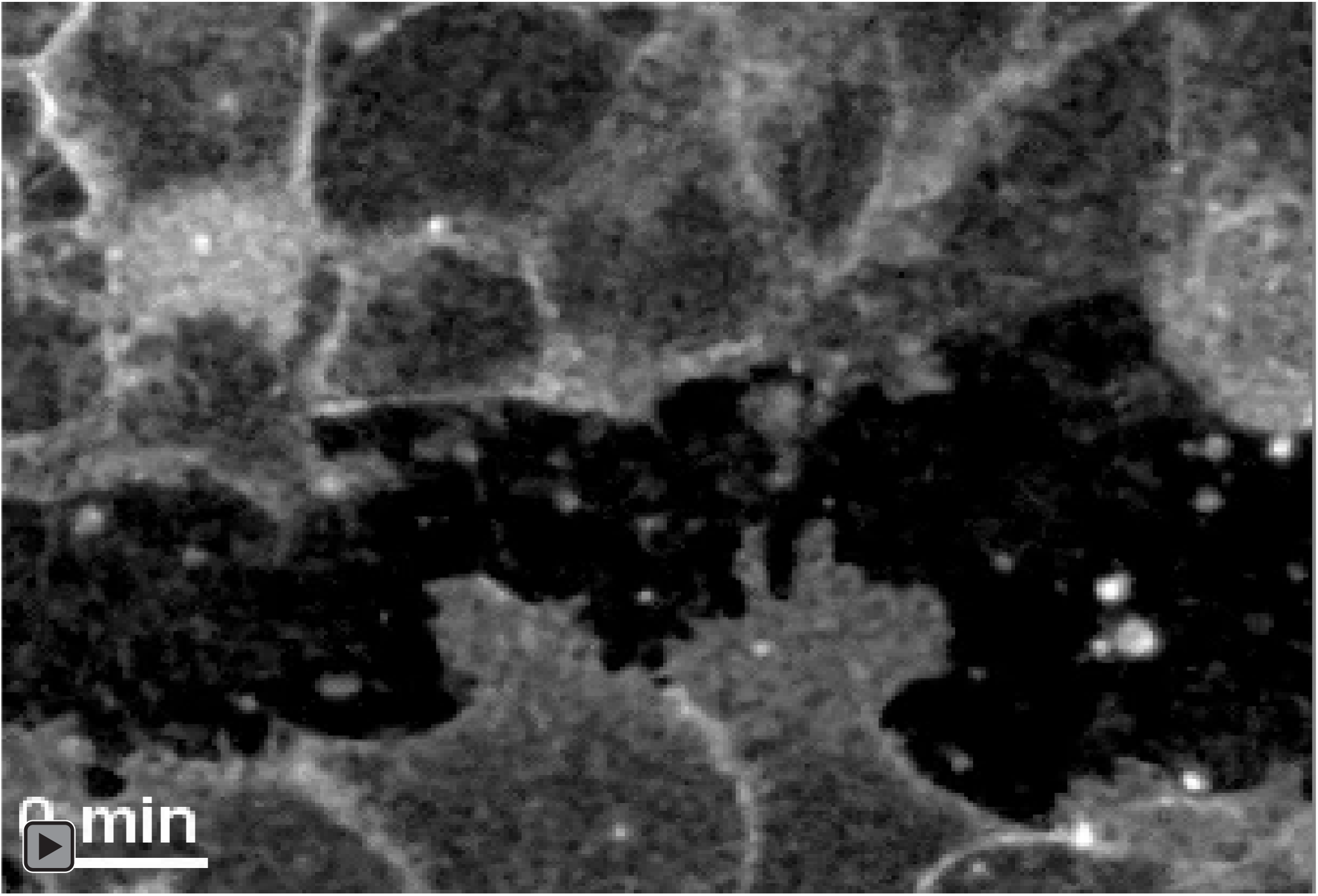
Mesendoderm closure of ΔE-cdh3 expressing embryo. Confocal time-lapse sequence of mesendoderm closure expressing ΔE-cdh3 and membrane-targeted GFP. Data were acquired every 15 seconds. Scale bar is 20 µm.

**Movie 4.**
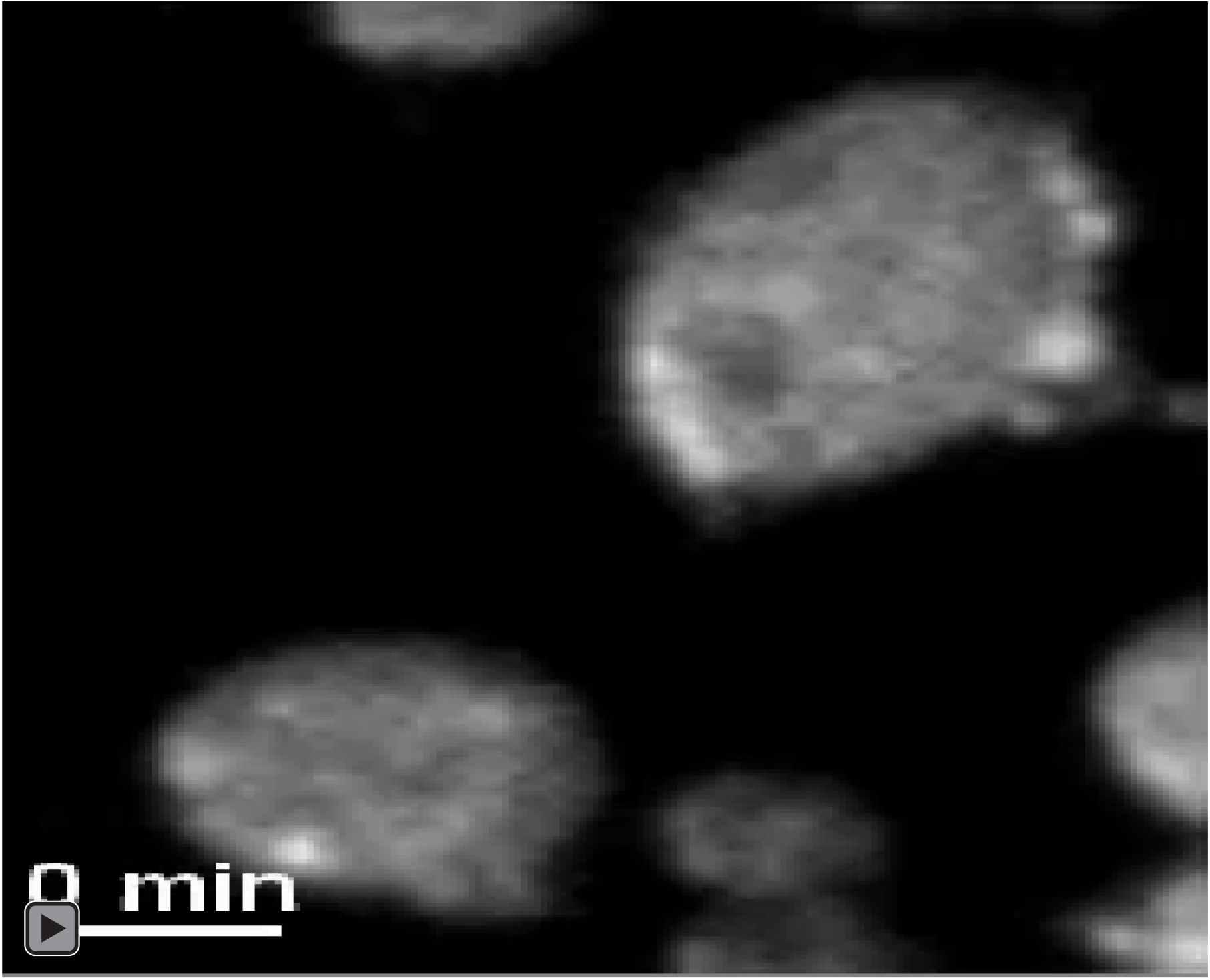
Single-cell CIL of ΔE-cdh3 expressing mesendoderm. Time-lapse sequence of single cells expressing ΔE-cdh3 observed using brightfield illumination on an inverted compound microscope. The time interval is 30 seconds. Scale bar is 20 µm.

**Movie 5.**
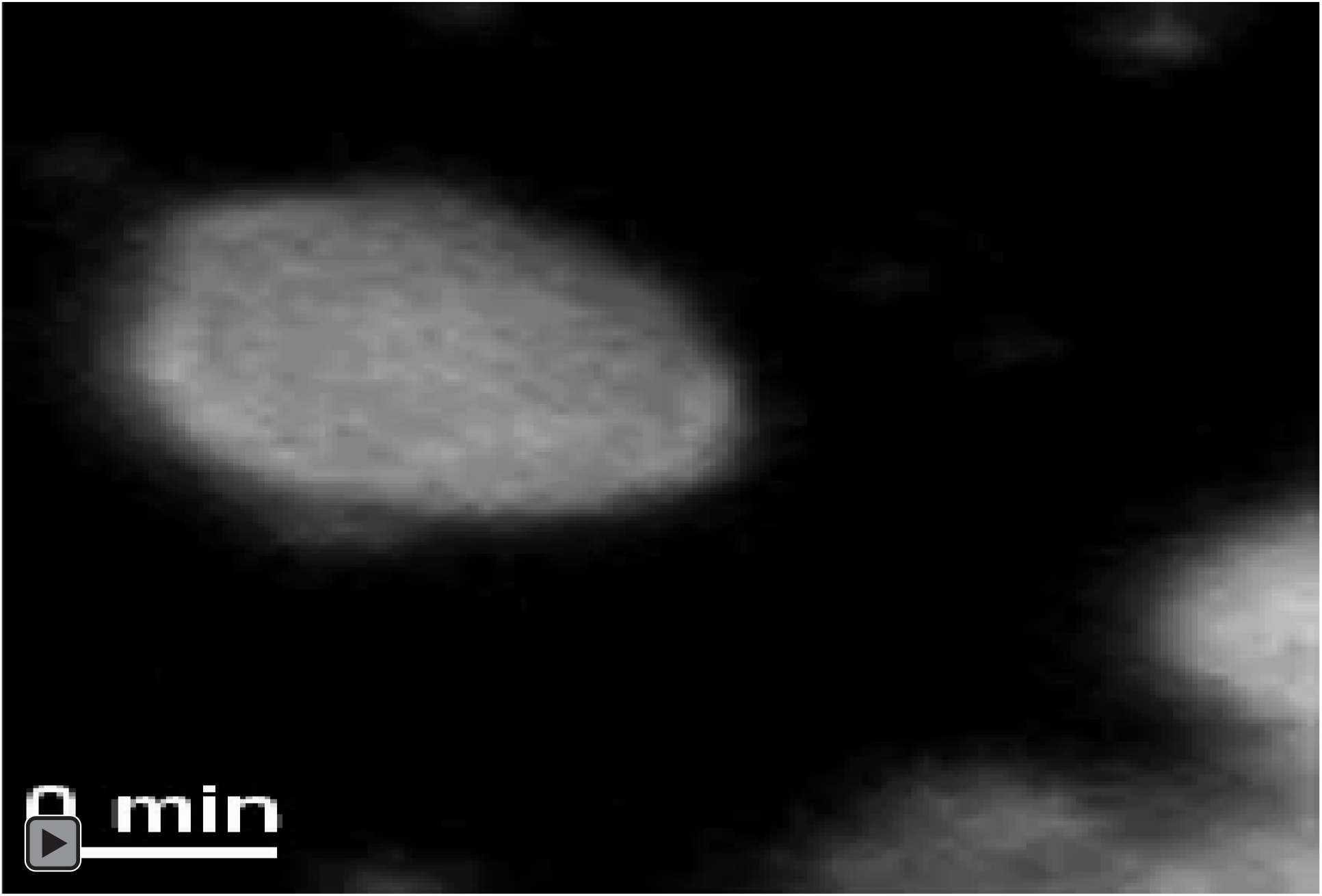
Single-cell CIL of ΔC-cdh3 expressing mesendoderm. Time-lapse sequence of single cells expressing ΔC-cdh3 observed using brightfield illumination on an inverted compound microscope. The time interval is 30 seconds. Scale bar is 20 µm.

**Movie 6.**
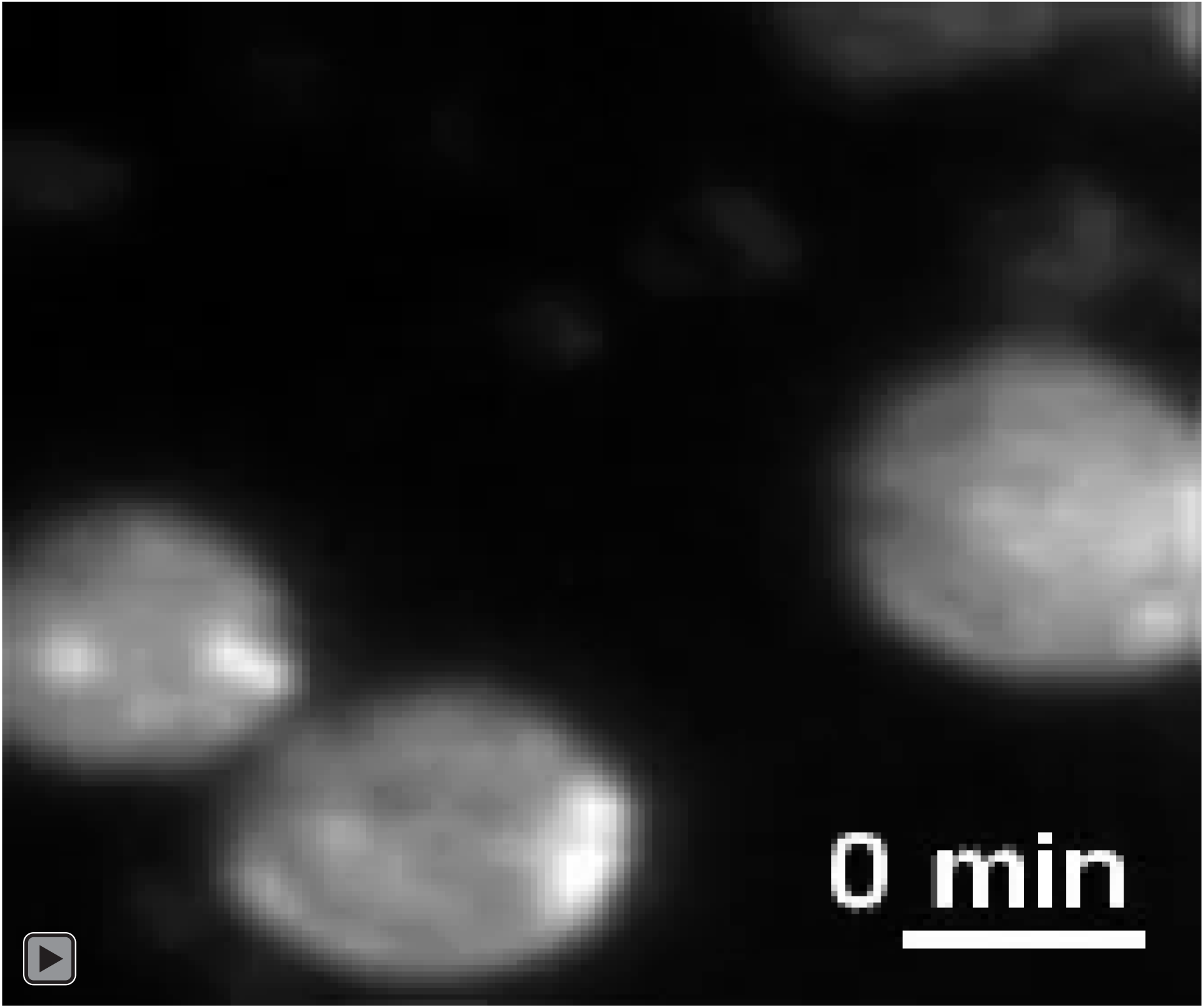
Single-cell CIL of ΔE-cdh3 + ΔC-cdh3 co-expressing mesendoderm. Time-lapse sequence of single cells co-expressing ΔE-cdh3 and ΔC-cdh3 observed using brightfield illumination on an inverted compound microscope. The time interval is 30 seconds. Scale bar is 20 µm.

**Movie 7.**
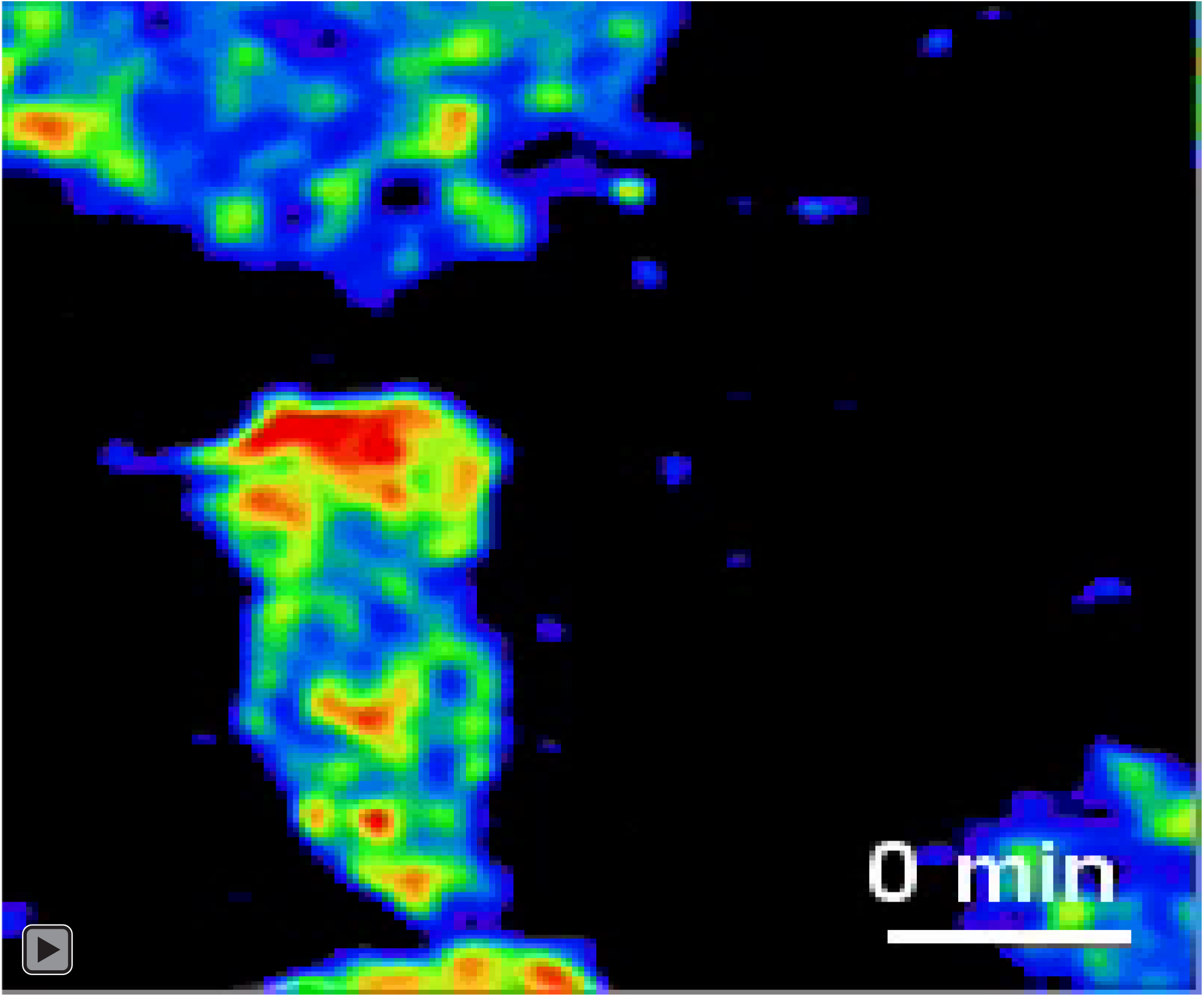
Subcellular Rac1 activity of wild-type mesendodermal cell during a cell-cell collision. Confocal time-lapse sequence of wild-type single-cell CIL showing Rac1 activity before, during, and after the cell-cell collision. Data were acquired every 15 seconds with a 63× lens and an inverted confocal microscope. Scale bar is 20 µm.

**Movie 8.**
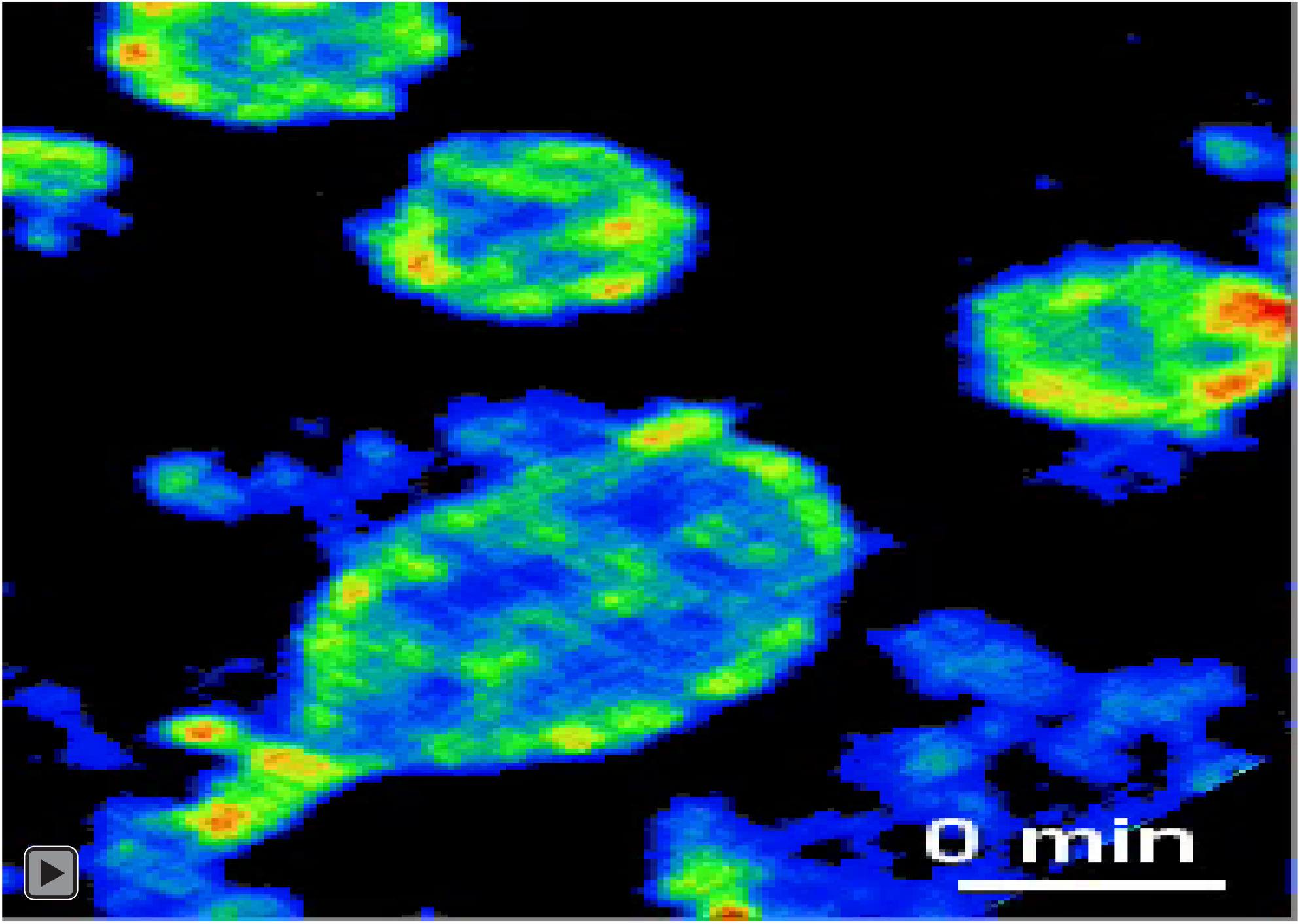
Subcellular Rac1 activity of ΔE-cdh3 mesendodermal cell during a cell-cell collision. Confocal time-lapse sequence of ΔE-cdh3 single cell CIL showing Rac1 activity before, during, and after the cell-cell collision. Data were acquired at every 15 seconds. Scale bar is 20 µm.

**Movie 9.**
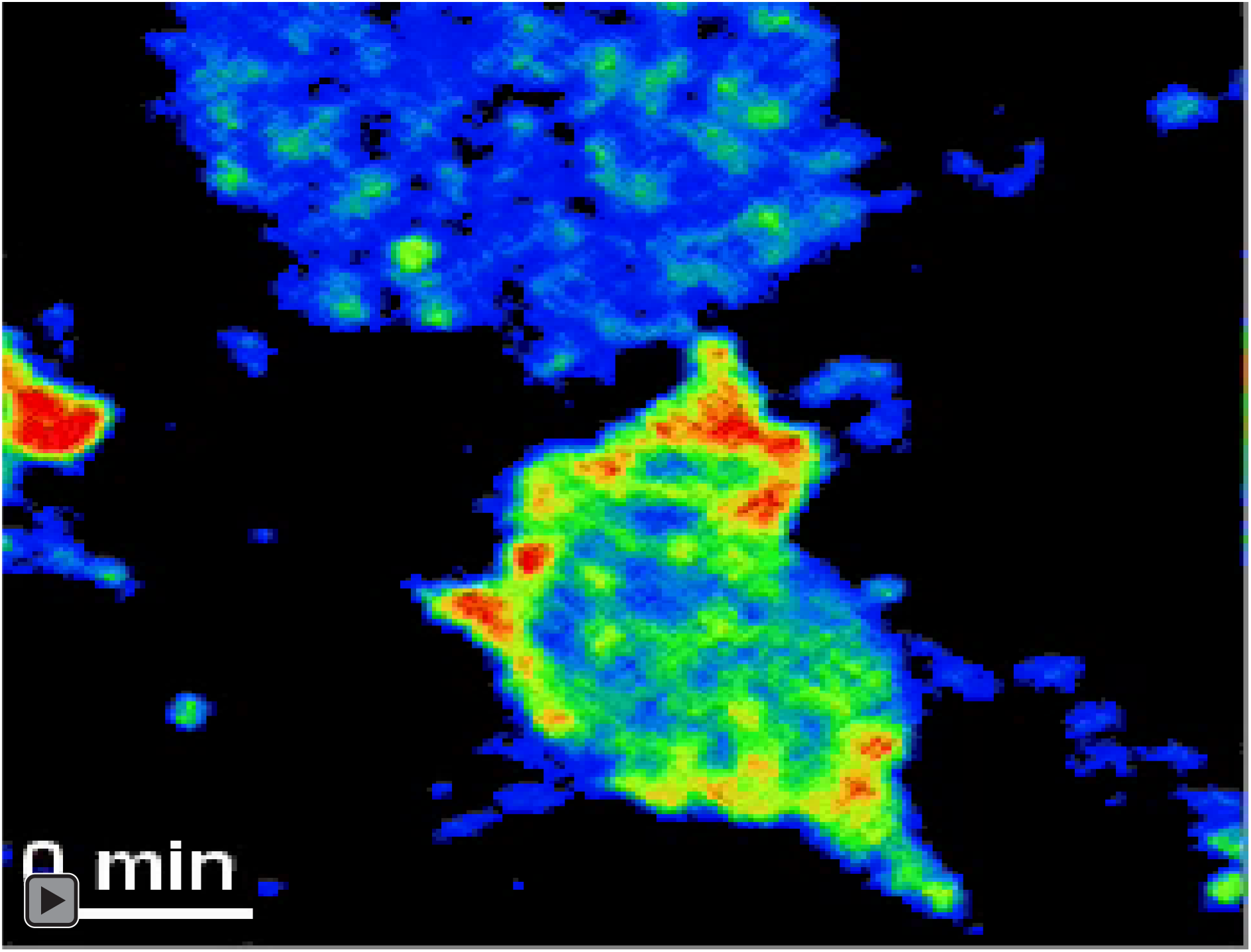
Subcellular Rac1 activity of ΔC-cdh3 mesendodermal cell during cell-cell collision. Time-lapse movie of ΔC-cdh3 single-cell CIL showing Rac1 activity before, during, and after the cell-cell collision. Data were acquired every 15 seconds. Scale bar is 20 µm.

**Table S1.** Mean angle and incidence of single-cell CIL.

| Sample | N | mean<br>(degree) | circular p | decision |
| --- | --- | --- | --- | --- |
| WT | 46 | 122.5 | < 0.05 | normal CIL |
| $\Delta E$-cdh3 | 37 | 7.6 | < 0.001 | deficient |
| $\Delta C$-cdh3 | 40 | -179.8 | < 0.01 | normal |
| $\Delta E$-cdh3 + $\Delta C$-cdh3 | 47 | -113.5 | 0.07 | partial rescue |
| $\Delta E$-cdh3 + FL-cdh3 | 37 | -138.9 | 0.69 | partial rescue |
| $\Delta E$-cdh3 + caRac | 42 | -167.8 | 0.43 | partial rescue |
| dnRac + RacMO | 44 | 9 | 0.08 | partial deficient |
| WT in 0.3 mM NSC23766 | 49 | -11.5 | 0.17 | partial deficient |
| $\Delta E$-cdh3 in 20 nM paclitaxel | 49 | -25 | 0.71 | partial rescue |
| $\Delta E$-cdh3 in 20 nM jasplakinolide | 37 | -22.9 | 0.09 | partial rescue |
| WT in 20 nM nocodazole | 38 | 30.7 | 0.19 | partial deficient |
| WT in 20 nM cytochalasin D | 42 | 7.1 | 0.87 | partial deficient |
| $\Delta E$-cdh3 to WT | 32 | -6.7 | < 0.001 | deficient |
| WT to $\Delta E$-cdh3 | 29 | 143.2 | < 0.001 | normal |
| WT to cdh3-beads | 41 | -1.6 | < 0.001 | deficient |

The number of samples, mean angle, the p-value of Rayleigh test, and classification of contact inhibition of locomotion (CIL) are shown. Mean angle was measured with respect to the incident angle of the approaching cell. Normal CIL was defined in collisions, where the mean angle was significantly greater than $120^\circ$ ( $p < 0.05$ ). Cases of deficient CIL occur where the mean angle was significantly less than $60^\circ$ ( $p < 0.05$ ). Other cases are qualified as partial rescue or partial deficiencies in CIL.

**Table S2.** Persistence (directionality) of single migrating cells.

| Conditions | N | directionality | SE | p-value of paired t-test<br>(for sample) |
| --- | --- | --- | --- | --- |
| WT | 54 | 0.47 | 0.0239 |  |
| $\Delta E$-cdh3 | 43 | 0.6 | 0.0313 | < 0.01 (WT) |
| $\Delta C$-cdh3 | 35 | 0.39 | 0.0327 | < 0.01 (WT) |
| $\Delta E$-cdh3 + $\Delta C$-cdh3 | 47 | 0.47 | 0.0173 | < 0.001 ($\Delta E$-cdh3)<br>< 0.001 ($\Delta C$-cdh3) |
| $\Delta E$-cdh3 + FL-cdh3 | 37 | 0.46 | 0.0295 | < 0.01 ($\Delta E$-cdh3) |
| $\Delta C$-cdh3 + FL-cdh3 | 36 | 0.43 | 0.032 | < 0.05 ($\Delta C$-cdh3) |
| $\Delta E$-cdh3 + caRac | 37 | 0.46 | 0.027 | < 0.01 ($\Delta E$-cdh3) |
| $\Delta C$-cdh3 + dnRac | 37 | 0.51 | 0.0282 | < 0.001 ($\Delta C$-cdh3) |
| caRac | 41 | 0.38 | 0.0314 | < 0.05 (WT) |
| dnRac + RacMO | 37 | 0.61 | 0.0276 | < 0.001 (WT) |
| $\Delta C$-cdh3 in 20 nM<br>NSC23766 | 49 | 0.56 | 0.0248 | < 0.001 ($\Delta C$-cdh3) |
| $\Delta E$-cdh3 in 20 nM<br>Pacritaxel | 52 | 0.39 | 0.0186 | < 0.001 ($\Delta E$-cdh3) |
| $\Delta E$-cdh3 in 20 nM<br>Jasplakinolide | 44 | 0.41 | 0.024 | < 0.001 ($\Delta E$-cdh3) |
| WT in 20 nM<br>Nocodazole | 51 | 0.53 | 0.0221 | < 0.05 (WT) |
| WT in 20 nM<br>Cytochalasin D | 37 | 0.55 | 0.0283 | < 0.01 (WT) |

The number of samples, directionality, standard error, and p-value of paired t-test for each sample in parentheses are shown. Values were collected by tracking single migratory cells for 1 hour without collision.

## References

Abercrombie, M. (1970). Contact inhibition in tissue culture. In Vitro 6, 128–142.

Abercrombie, M. and Heaysman, J. E. (1953). Observations on the social behaviour of cells in tissue culture. I. Speed of movement of chick heart fibroblasts in relation to their mutual contacts. Exp Cell Res 5, 111–131.

Abercrombie, M. and Heaysman, J. E. (1954). Observations on the social behaviour of cells in tissue culture. II. Monolayering of fibroblasts. Exp Cell Res 6, 293–306.

Anastasiadis, P. Z. and Reynolds, A. B. (2001). Regulation of Rho GTPases by p120-catenin. Curr Opin Cell Biol 13, 604–610.

Astin, J. W., Batson, J., Kadir, S., Charlet, J., Persad, R. A., Gillatt, D., Oxley, J. D. and Nobes, C. D. (2010). Competition amongst Eph receptors regulates contact inhibition of locomotion and invasiveness in prostate cancer cells. Nat Cell Biol 12, 1194–1204.

Baronsky, T., Dzementsei, A., Oelkers, M., Melchert, J., Pieler, T. and Janshoff, A. (2016). Reduction in E-cadherin expression fosters migration of Xenopus laevis primordial germ cells. Integr Biol (Camb) 8, 349–358.

Becker, S. F., Mayor, R. and Kashef, J. (2013). Cadherin-11 mediates contact inhibition of locomotion during Xenopus neural crest cell migration. PLoS One 8, e85717.

Carmona-Fontaine, C., Matthews, H. K., Kuriyama, S., Moreno, M., Dunn, G. A., Parsons, M., Stern, C. D. and Mayor, R. (2008). Contact inhibition of locomotion in vivo controls neural crest directional migration. Nature 456, 957–961.

Chappuis-Flament, S., Wong, E., Hicks, L. D., Kay, C. M. and Gumbiner, B. M. (2001). Multiple cadherin extracellular repeats mediate homophilic binding and adhesion. J Cell Biol 154, 231–243.

Choi, Y. S. and Gumbiner, B. (1989). Expression of cell adhesion molecule E-cadherin in Xenopus embryos begins at gastrulation and predominates in the ectoderm. J Cell Biol 108, 2449–2458.

Davidson, L. A., Hoffstrom, B. G., Keller, R. and DeSimone, D. W. (2002). Mesendoderm extension and mantle closure in *Xenopus laevis* gastrulation: combined roles for integrin alpha5beta1, fibronectin, and tissue geometry. Dev Biol 242, 109–129.

Davidson, L. A., Keller, R. and DeSimone, D. (2004). Patterning and tissue movements in a novel explant preparation of the marginal zone of Xenopus laevis. Gene Expr Patterns 4, 457–466.

Detrick, R. J., Dickey, D. and Kintner, C. R. (1990). The effects of N-cadherin misexpression on morphogenesis in Xenopus embryos. Neuron 4, 493–506.

Du, W., Liu, X., Fan, G., Zhao, X., Sun, Y., Wang, T., Zhao, R., Wang, G., Zhao, C., Zhu, Y., et al. (2014). From cell membrane to the nucleus: an emerging role of E-cadherin in gene transcriptional regulation. J Cell Mol Med 18, 1712–1719.

Dunn, G. A. and Paddock, S. W. (1982). Analysing the motile behaviour of cells: a general approach with special reference to pairs of cells in collision. Philos Trans R Soc Lond B Biol Sci 299, 147–157.

Fang, X., Ji, H., Kim, S. W., Park, J. I., Vaught, T. G., Anastasiadis, P. Z., Ciesiolka, M. and McCrea, P. D. (2004). Vertebrate development requires ARVCF and p120 catenins and their interplay with RhoA and Rac. J Cell Biol 165, 87–98.

Franz, C. M. and Ridley, A. J. (2004). p120 catenin associates with microtubules: inverse relationship between microtubule binding and Rho GTPase regulation. J Biol Chem 279, 6588–6594.

Gao, Y., Dickerson, J. B., Guo, F., Zheng, J. and Zheng, Y. (2004). Rational design and characterization of a Rac GTPase-specific small molecule inhibitor. Proceedings of the National Academy of Sciences of the United States of America 101, 7618–7623.

George, S. P., Chen, H., Conrad, J. C. and Khurana, S. (2013). Regulation of directional cell migration by membrane-induced actin bundling. J Cell Sci 126, 312–326.

Goodwin, M., Kovacs, E. M., Thoreson, M. A., Reynolds, A. B. and Yap, A. S. (2003). Minimal mutation of the cytoplasmic tail inhibits the ability of E-cadherin to activate Rac but not phosphatidylinositol 3-kinase: direct evidence of a role for cadherin-activated Rac signaling in adhesion and contact formation. J Biol Chem 278, 20533–20539.

Gottardi, C. J., Wong, E. and Gumbiner, B. M. (2001). E-cadherin suppresses cellular transformation by inhibiting beta-catenin signaling in an adhesion-independent manner. J Cell Biol 153, 1049–1060.

Habas, R., Dawid, I. B. and He, X. (2003). Coactivation of Rac and Rho by Wnt/Frizzled signaling is required for vertebrate gastrulation. Genes Dev 17, 295–309.

Hara, Y., Nagayama, K., Yamamoto, T. S., Matsumoto, T., Suzuki, M. and Ueno, N. (2013). Directional migration of leading-edge mesoderm generates physical forces: Implication in Xenopus notochord formation during gastrulation. Dev Biol 382, 482–495.

Huang, C., Kratzer, M. C., Wedlich, D. and Kashef, J. (2016). E-cadherin is required for cranial neural crest migration in Xenopus laevis. Dev Biol 411, 159–171.

Itoh, R. E., Kurokawa, K., Ohba, Y., Yoshizaki, H., Mochizuki, N. and Matsuda, M. (2002). Activation of rac and cdc42 video imaged by fluorescent resonance energy transfer-based single-molecule probes in the membrane of living cells. Mol Cell Biol 22, 6582–6591.

Kardash, E., Reichman-Fried, M., Maitre, J. L., Boldajipour, B., Papusheva, E., Messerschmidt, E. M., Heisenberg, C. P. and Raz, E. (2010). A role for Rho GTPases and cell-cell adhesion in single-cell motility in vivo. Nat Cell Biol 12, 47–53; sup pp 41-11.

Kaverina, I. and Straube, A. (2011). Regulation of cell migration by dynamic microtubules. Semin Cell Dev Biol 22, 968–974.

Kim, H. Y. and Davidson, L. A. (2011). Punctuated actin contractions during convergent extension and their permissive regulation by the non-canonical Wnt-signaling pathway. J Cell Sci 124, 635–646.

Kintner, C. (1992). Regulation of embryonic cell adhesion by the cadherin cytoplasmic domain. Cell 69, 225–236.

Kowalczyk, A. P. and Reynolds, A. B. (2004). Protecting your tail: regulation of cadherin degradation by p120-catenin. Curr Opin Cell Biol 16, 522–527.

Krause, M. and Gautreau, A. (2014). Steering cell migration: lamellipodium dynamics and the regulation of directional persistence. Nat Rev Mol Cell Biol 15, 577–590.

Kraynov, V. S., Chamberlain, C., Bokoch, G. M., Schwartz, M. A., Slabaugh, S. and Hahn, K. M. (2000). Localized Rac activation dynamics visualized in living cells. Science 290, 333–337.

Kumper, S. and Ridley, A. J. (2010). p120ctn and P-cadherin but not E-cadherin regulate cell motility and invasion of DU145 prostate cancer cells. PLoS One 5, e11801.

Kurth, T., Fesenko, I. V., Schneider, S., Munchberg, F. E., Joos, T. O., Spieker, T. P. and Hausen, P. (1999). Immunocytochemical studies of the interactions of cadherins and catenins in the early Xenopus embryo. Dev Dyn 215, 155–169.

Lee, C. H. and Gumbiner, B. M. (1995). Disruption of gastrulation movements in Xenopus by a dominant-negative mutant for C-cadherin. Dev Biol 171, 363–373.

Levine, E., Lee, C. H., Kintner, C. and Gumbiner, B. M. (1994). Selective disruption of E-cadherin function in early Xenopus embryos by a dominant negative mutant. Development 120, 901–909.

Liao, Z., Cao, C., Wang, J., Huxley, V. H., Baker, O., Weisman, G. A. and Erb, L. (2014). The P2Y Receptor Interacts with VE-Cadherin and VEGF Receptor-2 to Regulate Rac1 Activity in Endothelial Cells. J Biomed Sci Eng 7, 1105–1121.

Liu, W. F., Nelson, C. M., Pirone, D. M. and Chen, C. S. (2006). E-cadherin engagement stimulates proliferation via Rac1. J Cell Biol 173, 431–441.

Machacek, M., Hodgson, L., Welch, C., Elliott, H., Pertz, O., Nalbant, P., Abell, A., Johnson, G. L., Hahn, K. M. and Danuser, G. (2009). Coordination of Rho GTPase activities during cell protrusion. Nature 461, 99–103.

Mateus, A. R., Seruca, R., Machado, J. C., Keller, G., Oliveira, M. J., Suriano, G. and Luber, B. (2007). EGFR regulates RhoA-GTP dependent cell motility in E-cadherin mutant cells. Hum Mol Genet 16, 1639–1647.

Matthews, H. K., Marchant, L., Carmona-Fontaine, C., Kuriyama, S., Larrain, J., Holt, M. R., Parsons, M. and Mayor, R. (2008). Directional migration of neural crest cells in vivo is regulated by Syndecan-4/Rac1 and non-canonical Wnt signaling/RhoA. Development 135, 1771–1780.

Mayor, R. and Carmona-Fontaine, C. (2010). Keeping in touch with contact inhibition of locomotion. Trends Cell Biol 20, 319–328.

Monypenny, J., Zicha, D., Higashida, C., Oceguera-Yanez, F., Narumiya, S. and Watanabe, N. (2009). Cdc42 and Rac family GTPases regulate mode and speed but not direction of primary fibroblast migration during platelet-derived growth factor-dependent chemotaxis. Mol Cell Biol 29, 2730–2747.

Muller, H. A., Kuhl, M., Finnemann, S., Schneider, S., van der Poel, S. Z., Hausen, P. and Wedlich, D. (1994). Xenopus cadherins: the maternal pool comprises distinguishable members of the family. Mech Dev 47, 213–223.

Niessen, C. M. and Gottardi, C. J. (2008). Molecular components of the adherens junction. Biochim Biophys Acta 1778, 562–571.

Noren, N. K., Arthur, W. T. and Burridge, K. (2003). Cadherin engagement inhibits RhoA via p190RhoGAP. J Biol Chem 278, 13615–13618.

Noren, N. K., Niessen, C. M., Gumbiner, B. M. and Burridge, K. (2001). Cadherin engagement regulates Rho family GTPases. J Biol Chem 276, 33305–33308.

Padmanabhan, A., Ong, H. T. and Zaidel-Bar, R. (2017). Non-junctional E-Cadherin Clusters Regulate the Actomyosin Cortex in the C. elegans Zygote. Curr Biol 27, 103–112.

Pankov, R., Endo, Y., Even-Ram, S., Araki, M., Clark, K., Cukierman, E., Matsumoto, K. and Yamada, K. M. (2005). A Rac switch regulates random versus directionally persistent cell migration. J Cell Biol 170, 793–802.

Perez, T. D., Tamada, M., Sheetz, M. P. and Nelson, W. J. (2008). Immediate-early signaling induced by E-cadherin engagement and adhesion. J Biol Chem 283, 5014–5022.

Petrie, R. J., Doyle, A. D. and Yamada, K. M. (2009). Random versus directionally persistent cell migration. Nat Rev Mol Cell Biol 10, 538–549.

Plutoni, C., Bazellieres, E., Le Borgne-Rochet, M., Comunale, F., Brugues, A., Seveno, M., Planchon, D., Thuault, S., Morin, N., Bodin, S., et al. (2016). P-cadherin promotes collective cell migration via a Cdc42-mediated increase in mechanical forces. J Cell Biol 212, 199–217.

Rebman, J. K., Kirchoff, K. E. and Walsh, G. S. (2016). Cadherin-2 Is Required Cell Autonomously for Collective Migration of Facial Branchiomotor Neurons. PLoS One 11, e0164433.

Roca-Cusachs, P., Sunyer, R. and Trepat, X. (2013). Mechanical guidance of cell migration: lessons from chemotaxis. Curr Opin Cell Biol 25, 543–549.

Rottner, K., Hall, A. and Small, J. V. (1999). Interplay between Rac and Rho in the control of substrate contact dynamics. Curr Biol 9, 640–648.

Scarpa, E., Szabo, A., Bibonne, A., Theveneau, E., Parsons, M. and Mayor, R. (2015). Cadherin Switch during EMT in Neural Crest Cells Leads to Contact Inhibition of Locomotion via Repolarization of Forces. Developmental cell 34, 421–434.

Sive, H. L., Grainger, R. M. and Harland, R. M. (2000). Early development of Xenopus laevis: a laboratory manual. Cold Spring Harbor, N.Y.: Cold Spring Harbor Laboratory Press.

Theveneau, E., Steventon, B., Scarpa, E., Garcia, S., Trepat, X., Streit, A. and Mayor, R. (2013). Chase-and-run between adjacent cell populations promotes directional collective migration. Nat Cell Biol 15, 763–772.

Troxell, M. L., Chen, Y.-T., Cobb, N., Nelson, W. J. and Marrs, J. A. (1999). Cadherin function in junctional complex rearrangement and posttranslational control of cadherin expression. American Journal of Physiology-Cell Physiology 276, C404–C418.

Valls, G., Codina, M., Miller, R. K., Del Valle-Perez, B., Vinyoles, M., Caelles, C., McCrea, P. D., Garcia de Herreros, A. and Dunach, M. (2012). Upon Wnt stimulation, Rac1 activation requires Rac1 and Vav2 binding to p120-catenin. J Cell Sci 125, 5288–5301.

Vincent, S., Brouns, M., Hart, M. J. and Settleman, J. (1998). Evidence for distinct mechanisms of transition state stabilization of GTPases by fluoride. Proc Natl Acad Sci U S A 95, 2210–2215.

Weber, G. F., Bjerke, M. A. and DeSimone, D. W. (2012). A mechanoresponsive cadherin-keratin complex directs polarized protrusive behavior and collective cell migration. Developmental cell 22, 104–115.

Winklbauer, R. (2012). Cadherin function during Xenopus gastrulation. Subcell Biochem 60, 301–320.

Winklbauer, R. and Keller, R. E. (1996). Fibronectin, mesoderm migration, and gastrulation in Xenopus. Dev Biol 177, 413–426.

Winklbauer, R. and Nagel, M. (1991). Directional mesoderm cell migration in the Xenopus gastrula. Dev Biol 148, 573–589.

Winklbauer, R., Selchow, A., Nagel, M. and Angres, B. (1992). Cell interaction and its role in mesoderm cell migration during Xenopus gastrulation. Dev Dyn 195, 290–302.

Wittmann, T., Bokoch, G. M. and Waterman-Storer, C. M. (2003). Regulation of leading edge microtubule and actin dynamics downstream of Rac1. J Cell Biol 161, 845–851.

Yamao, M., Naoki, H., Kunida, K., Aoki, K., Matsuda, M. and Ishii, S. (2015). Distinct predictive performance of Rac1 and Cdc42 in cell migration. Sci Rep 5, 17527.

Zar, J. H. (1999). Biostatistical analysis (4th edn). Upper Saddler River, NJ: Prentice Hall.

Zhang, B., Chernoff, J. and Zheng, Y. (1998). Interaction of Rac1 with GTPase-activating proteins and putative effectors. A comparison with Cdc42 and RhoA. J Biol Chem 273, 8776–8782.

## Reference

1. Kardash, E. et al. A role for Rho GTPases and cell-cell adhesion in single-cell motility in vivo. Nat Cell Biol 12, 47–53; sup pp 41-11, doi:10.1038/ncb2003 ncb2003 [pii] (2010).

